# Predicting Saturation Concentrations of Phase-Separating Proteins via Thermodynamic Integration

**DOI:** 10.1101/2025.05.09.653068

**Authors:** Eduardo Pedraza, Andrés R. Tejedor, Alejandro Feito, Francisco Gámez, Rosana Collepardo-Guevara, Eduardo Sanz, Jorge R. Espinosa

## Abstract

Phase separation of proteins and nucleic acids into biomolecular condensates contributes to the regulation of cellular compartmentalisation in membrane-less environments. A key parameter controlling the onset of biomolecular condensate formation via liquid—liquid phase separation is the saturation concentration (*C*_*sat*_)— the threshold concentration above which condensation takes place. While measuring *C*_*sat*_ for protein solutions *in vitro* is experimentally accessible, determining this quantity in simulations remains challenging due to the extremely low equilibrium concentrations at which many proteins phase separate. This occurs because the gold standard in simulations consists on combining a residue-resolution coarse-grained model with the Direct Coexistence simulation method, which yields poor estimates of the equilibrium concentrations of the dilute phase due to lack of statistics. In this work, we present two independent thermodynamic integration (TI) schemes which, when combined with Direct Coexistence simulations, enable accurate calculation of saturation concentrations and phase diagrams—facilitating direct comparison with experimental measurements across a wide range of conditions. Our methods, combined with the Mpipi-Recharged residue-resolution coarse-grained model, accurately estimate *C*_*sat*_ for a wide range of intrinsically disordered and multi-domain proteins, including disease-associated RNA- and DNA-binding proteins involved in the formation of stress granules and P granules, as well as engineered mutants of hnRNPA1. Furthermore, we compare our TI methods against a computationally efficient machine-learning predictor trained to estimate saturation concentrations at sub-physiological temperatures. While both approaches yield realistic predictions, explicit molecular dynamics simulations enable the calculation of complete phase diagrams and provide insight into the molecular mechanisms and interactions driving phase-separation. Overall, our approach offers a robust, physically grounded framework for improving and validating coarse-grained models of biomolecular phase behaviour, effectively bridging the gap between simulation and experiment.

## INTRODUCTION

Liquid-liquid phase separation (LLPS) is a fundamental mechanism used by cells to spatiotemporally compartmentalise their material and perform wide-ranging biological functions^1–4^. LLPS of proteins and nucleic acids has emerged as a central area of research in molecular biology, biophysics, and computational biology due to its large implications in health and disease^5–9^. Through this mechanism, biomolecules including intrinsically disordered proteins, multi-domain proteins, RNAs, or DNAs spontaneously self-assemble into membraneless organelles, termed condensates, which consist of enriched phases in different types of biomolecules^8,10^. Bio-condensates are formed both in the cytosol—such as stress granules^11^, processing bodies^12,13^, or germ granules^14^—and in the nucleus, including the nucleoli^15^, Cajal bodies^16^, and paraspeckles^17^. These structures enable spatiotemporal control over biomolecular organization and regulate functions such as RNA metabolism^18^, signal transduction^19^, and stress responses^20^, among many others^21–23^. Understanding LLPS is crucial not only for elucidating the mechanisms of cellular compartmentalisation, but also for shedding light on aberrant liquid-to-solid transitions of condensates into toxic solid-like assemblies ^24,25^—a phenomenon linked to multiple neurodegenerative disorders such as amyotrophic lateral sclerosis (ALS)^6^, frontotemporal dementia (FTD)^26,27^, Parkinson^28^, and Alzheimer’s disease^29^. Therefore, determining the physicochemical properties of the biomolecules regulating LLPS is vital for dissecting the thermodynamic and molecular mechanisms governing the dynamic formation and dissolution of condensates.

Multivalent proteins with intrinsically disordered regions (IDRs) act as key drivers of LLPS by locally establishing homotypic and heterotypic interactions with cognate biomolecules such as RNAs, DNAs, or other proteins^30–32^. Archetypal proteins inducing condensate formation include RNA-binding proteins such as FUS^6,33^, hnRNPA1^34–36^, LAF-1^37^, or Y-box1^38^, and DNA-binding proteins such as TDP-43^39–41^ or HP1^42,43^. Subtle variations of temperature, pH, salt concentration, or the presence of other biomolecules like sorbitol or glucose in the cell environment lead to drastic changes in the phase separation propensity of proteins^30,32,44–47^. Thus, determining the critical concentration to undergo phase-separation—usually termed as saturation concentration (*C*_*sat*_)—is essential to understand and potentially modulate the propensity of biomolecules to form condensates^11,48–52^.

*In vitro* assays have been shown to be the most efficient experimental framework to explore biomolecular condensation under specific well-controlled conditions^30,44,53–56^. Importantly, advanced super-resolution microscopy and fluorescence spectroscopies are capable of accessing the optimal length and timescales to investigate phase separation and coalescence times, including stimulated emission depletion (STED) microscopy^57,58^, Raman spectroscopy^59,60^, fluorescence correlation spectroscopy (FCS)^30,45,61^ and fluorescence recovery after photobleaching (FRAP)^30,46,62–64^. In particular, the protein saturation concentration required to form condensates is a key observable quantified in these experiments under specific conditions of pH, temperature, and salt concentration. Recent advances in experimental techniques have allowed the systematic evaluation of accurate *C*_*sat*_ measurements as a function of temperature^65–67^. Nevertheless, evaluating the protein concentration within the coexisting condensates, giving rise to full phase diagrams in the temperature–concentration plane, remains challenging ^14,65^. However, while experimentally measuring densities or concentrations inside the condensates remains elusive, performing it through computer simulations is relatively straightforward^68–71^. In contrast, determining the concentration of biomolecules in the dilute phase is highly challenging and requires vast sampling^72,73^, being most of the times computationally unfeasible due to the extremely low protein concentrations in such phase (*i*.*e*., 1-50 *µ*M^65,67^). Hence, directly comparing experimental and computational phase diagrams becomes a non-trivial task.

In the last decade, computational advances have enabled the characterisation of biomolecular phase separation with unprecedented accuracy and efficiency^69,71,74,75^. Continuous cross-validation and combination of experimental results, theoretical approaches, and molecular dynamics (MD) simulations have become fundamental to deepen our understanding of LLPS^69,76,77^. This multidisciplinary and integrative approach has been crucial for elucidating the underlying molecular grammar in proteins and nucleic acids controlling intracellular organisation^78^. At the highest resolution, atomistic MD simulations have enabled the evaluation of the precise interactions between different amino acids—*i*.*e*., *π − π* stacking, cation-*π* interactions, hydrogen bonding, or electrostatic forces—governing phase-separation ^32,79–82^. At the other extreme, low-resolution coarse-grained (CG) models have delved into phase separation thermodynamics by simplifying the intermolecular interactions through approaches such as using multivalent patchy particles^83–85^, minimal polymer physics models^79,86,87^, or by means of lattice-based simulations^88–91^. Furthermore, bioinformatic tools based on sequence characteristics such as IDR composition, secondary structural propensity, and charge patterning^92–94^, including PLAAC^92^, catGranule^95^, or PScore^96^ softwares have pushed forward our understanding of biomolecular phase separation by predicting LLPS propensity. Complementarily, recent advances in machine-learning (ML) and big data algorithms have revolutionised LLPS predictors by leveraging large datasets including structural and sequence information, from both experimental and computational libraries, to identify residue motifs and biophysical parameters associated with phase separation in diverse protein families and mixtures^97–102^. In that sense, ML predictors offer a scalable solution for proteome-wide LLPS analysis, and they are particularly optimal to carry out fast-screening analysis of hundreds or even thousands of protein sequences^71,103–105^. Nonetheless, ML predictors present two major drawbacks: 1) they usually rely on former coarse-grained high-resolution models and bioinformatic schemes that can bias their predictions^71,103^; and 2) their ‘black-box’ intrinsic nature hampers the rationalisation of the molecular mechanisms and intermolecular interactions controlling LLPS.

Chemically-accurate residue-resolution models lie at the interface between accurate physicochemical realism and computational efficiency for simulating biomolecular condensates^69,70^. These models portray a protein as a multi-domain co-polymer whose beads represent amino acids with their own chemical identity and unique hydrophobic and electrostatic interactions^70,77,91,106-108^. By integrating pair-specific parameters derived from atomistic simulations, experimental data, and bioinformatics, these models enhance the accuracy of predictions for single-molecule properties such as radii of gyration, and condensed phases, as critical solution temperatures or relative condensate viscosities, offering molecular and mechanistic rationalisation of the underlying forces behind LLPS^47,69,72,109,110^. Importantly, classical computational techniques from soft-matter physics can be incorporated in these models to harvest further information on the key physicochemical factors controlling condensate phase behaviour. For example, the implementation of the Green-Kubo relation and oscillatory shear simulations have recently allowed to characterise the viscoelastic properties of biomolecular condensates under equilibrium, and ageing conditions ^47,111–114^. In that respect, estimating the phase boundaries of a given biomolecular condensate—which are difficult to access for the dilute phase coexisting with the condensate^68,72^ using the gold standard method (Direct Coexistence (DC) simulations^74,115,116^)—could be approached from a statistical mechanics perspective using known thermodynamic relations.

In this work, we propose two different thermodynamic integration (TI) schemes to evaluate protein saturation concentrations of biomolecular condensates using sequence-dependent coarse-grained models in a straightforward manner. We employ the Mpipi-Recharged model^108^ combined with Direct Coexistence simulations and our TI approaches to accurately determine the coexistence protein concentration of the dilute phase across the binodal. Our two methods enable the calculation of extremely low saturation concentrations—as those experimentally found for biocondensates^14,30,65,67^—as a function of temperature based on quick DC simulations and bulk NVT calculations that evaluate protein condensate concentrations and internal energies. We validate our methods using both IDPs and multi-domain proteins, and we systematically apply TI calculations for a wide variety of phase separating proteins, such as FUS, LAF-1, hnRNPA1, TDP-43, G3BP1, DDX4, YBX1, and different constructs of them comparing our predictions with experimental *in vitro* measurements. Furthermore, we benchmark the predictions of the Mpipi-Recharged regarding the impact of different mutations in the low-complexity domain (LCD) of hnRNPA1 on the phase diagram. Finally, we compare saturation concentration predictions using the Mpipi-Recharged and our TI schemes against a ML predictor based on the CALVA-DOS2 model^71^ and *versus in vitro* measurements^65,67^. Taken together, our work proposes a robust framework to obtain saturation concentrations and directly validate modelling predictions against experimental datasets.

## METHODOLOGY

The saturation concentration is a fundamental magnitude to primarily characterise the ability of a biomolecular system to undergo phase-separation^30,48,50,65–67^. In a temperature–concentration phase diagram, the coexistence line for the dilute phase corresponds to the saturation concentration dependence with temperature. While evaluating *C*_*sat*_ either at near-critical solution temperatures or for molecules with low propensity to undergo phase-separation (*i*.*e*., with high *C*_*sat*_) is feasible through DC simulations^47,72,73^, quantifying *C*_*sat*_ for systems with extremely low coexisting densities is particularly challenging due to the scarce sampling in the dilute phase^47,72^. For that reason, here we propose two different thermodynamic integration routes, TI-*C*_*sat*_ and TI-*CC*, to accurately determine the saturation concentration from phase diagrams of biomolecular condensates evaluated via DC simulations. We employ the Mpipi-Recharged model—a residue-resolution CG model recently developed by us^68,69,108^—to simulate the formation of protein condensates via LLPS. Both methods proposed in this study enable calculating the coexistence density (or saturation concentration) of the dilute (protein-poor) phase at a low temperature, *T*_2_, from a coexistence point at a high temperature, *T*_1_, at which the densities of the condensed (protein-rich) and dilute phases (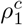and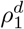, respectively) can be accurately obtained through density profiles in standard DC simulations (see Fig. 1a).

**FIG. 1.**
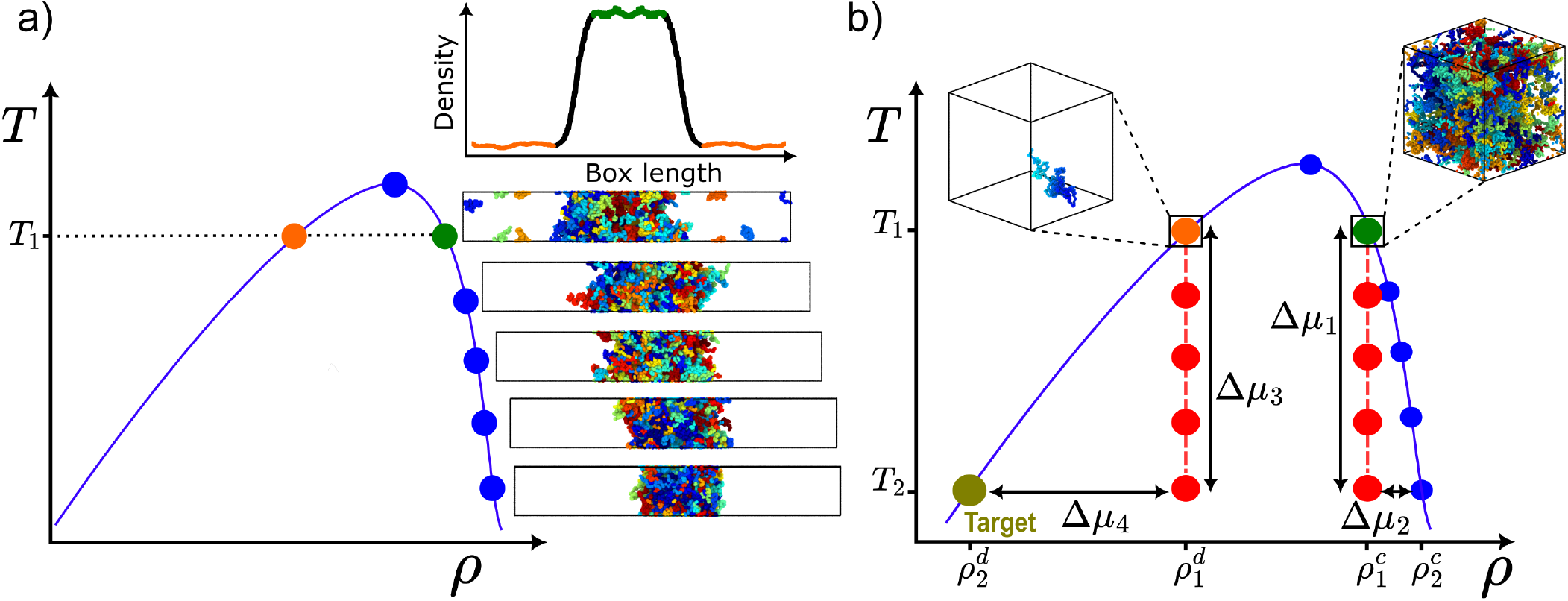
Overview of the TI-*C*_*sat*_ scheme. (a) Temperature–density (or concentration) phase diagram evaluated via Direct Coexistence simulations at different temperatures. The density of the coexisting phases is computed by applying a density profile along the long axis of the simulation box. Only at temperatures close to the critical one, the equilibrium density of the protein-poor liquid phase can be reliably measured. (b) Thermodynamic route across the phase diagram to compute the coexisting density (or saturation concentration) with a chemical potential 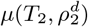 starting from the known densities 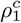 and 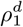 at temperature, *T*_1_. The red points represent *NV T* simulations at different temperatures, used to obtain Δ*µ*_1_ and Δ*µ*_3_ through thermodynamic integration, with the configurations for the dilute (left) and condensed (right) protein phases.

### A. The TI-C_sat_ method for computing saturation concentrations

The TI-*C*_*sat*_ method is based on the equality of chemical potentials at coexistence at both temperatures: 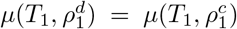 and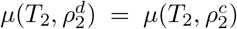. The following expression for 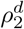 is obtained after connecting both coexistence points through a thermodynamic route sketched in Fig. 1b and explained in detail in the Appendix:

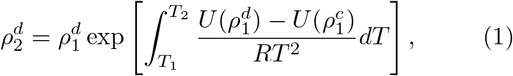

where and 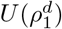 and 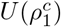 refer to the internal energy along the isochores at densities 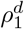 and 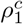, respectively. Hence, the target density 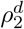 is calculated by evaluating 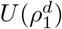 and 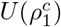 at different temperatures between *T*_2_ and *T*_1_ (as indicated by the red points in Fig. 1b). These calculations can be performed through standard *NV T* simulations of the bulk phases at their corresponding densities, 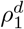 and 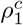, in temperature steps of 10-20K. Finally, the data are numerically integrated as written in Eq. (1), and the saturation concentration *C*_*sat*_ is straightforwardly calculated from 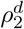 (*e*.*g*. if *ρ*_*d*_ is the number density and the volume is in liters, it is only required to divide over the Avogadro’s number).

### B. The Clausius-Clapeyron equation to determine C_sat_

In this alternative method, we assume the ideal gas approximation for the protein-poor liquid branch of the phase diagram. We exploit such approximation (also employed for the TI-*C*_*sat*_ as described in the Appendix) to calculate the target dilute density (or saturation concentration) through the Clausius-Clapeyron relation. We start from the general form of the Clapeyron equation^117–119^ given by:

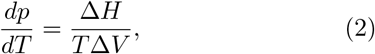

where Δ*H* and Δ*V* refer to the enthalpy difference and volume difference, respectively, between both coexisting phases in bulk conditions at a given temperature *T* and pressure *p*. Assuming the ideal gas equation, and given that the volume per protein within the condensate is negligible compared to the one in the dilute phase, we can integrate Eq. 2 to obtain:

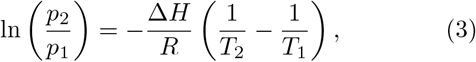

where *p*_1_ and *p*_2_ correspond to the equilibrium pressure at *T*_1_ and *T*_2_, respectively. The enthalpy difference between the coexistence phases can be approximated as the internal energy (*H ≈ U*) since the *pV* term in the enthalpy expression (*H* = *U* + *pV*) becomes negligible at equilibrium due to the practically zero pressure. Therefore, we can use the coexistence densities 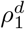 and 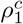 at near-critical conditions (*e*.*g*., orange and green circles in Fig. 1a estimated from DC simulations) and calculate the internal energy difference using NVT simulations in bulk conditions imposing the coexistence volume of each phase. Hence, from Eq. 3 we can obtain *p*_2_ at the target temperature *T*_2_ and subsequently calculate *C*_*sat*_ applying the ideal gas law equation to determine the molar volume associated with such coexistence pressure at *T*_2_:

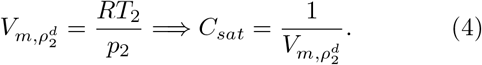

Importantly, this TI scheme assumes that Δ*H* (or Δ*U*) remains constant with temperature, but this approximation is only reasonable for moderate temperature intervals of 20-30 K. Thus, for accurately evaluating *C*_*sat*_ for |Δ*T*| *≳* 30*K*, we reassess the internal energy from NVT simulations at the bulk coexisting densities given by the calculated *C*_*sat*_ from the TI-*CC* approach at *T*_2_, and the condensed phase density from DC simulations. It is worth highlighting that to determine the internal energy of the dilute phase along different temperatures, it is sufficient to perform NVT simulations of a single molecule at each temperature in the dilute limit. Since the intermolecular interactions are negligible in the dilute limit, a sufficiently large simulation box can be used to isolate the molecule and compute the temperature-dependent internal energy contributions arising from intramolecular forces and the kinetic term.

## RESULTS AND DISCUSSION

### A. Validation of the thermodynamic integration schemes to compute protein condensate saturation concentrations

We first apply our two methods, TI-*C*_*sat*_ and TI-*CC*, for several archetypal proteins known to undergo LLPS. Specifically, we focus on Fused in Sarcoma (FUS)^33,50^, Y-box binding protein 1 (YBX1)^38^, and the arginine-glycine-rich region (RGG) of the Long Aspergillus-like Filament protein 1 (LAF-1-RGG)^37^ due to their distinct sequence composition, length, structure, and biological function. FUS is an RNA-binding protein composed of 526 amino acids, which includes two globular domains, and it is involved in different RNA metabolism processes and genome stability^6^. YBX1 is a protein formed by 324 amino acids that features a small globular domain, and acts as a recruiter of RNAs into exosomes and stress granules^38,120^. In contrast, LAF-1-RGG is an intrinsically disordered region of 176 amino acids which contributes to form cytoplasmic granules^37,121^. We use the Mpipi-Recharged model^108^ to simulate phase separation of FUS, YBX1, and LAF-1-RGG via DC simulations (please see Section SII of SM for further details) and compute their phase diagrams. The Mpipi-Recharged is a coarse-grained force field in which each amino acid is represented by a single bead with its own chemical identity.

Within this model, the intrinsically disordered regions are considered as fully flexible polymers, in which subsequent amino acids are connected by harmonic bonds, while the globular domains are treated as rigid bodies^108^ preserving the structures taken from the corresponding Protein Data Bank (PDB). The residue-residue interactions in the model consist of a combination of electrostatic and hydrophobic interactions^73,106^ implemented through a Yukawa potential and a Wang-Frenkel potential^122^, respectively (further technical details on the model can be found in Section SI of the SM). Importantly, the Yukawa potential permits the parameters to be finetuned depending on the specific charged residue compensating for the loss of explicit ions and water, as well as for the aggressive mapping of the many charges that an amino acid carries atomistically when being reduced to just one charge centered on its alpha carbon upon the coarse-graining. This model has been been successfully validated for reproducing single-protein radii of gyration, phase diagrams, and relative condensate viscosities of a relatively large set of protein sequences^68,69,108,123^.

In Fig. 2 we show the phase diagrams of FUS, YBX1 and LAF-1-RGG computed via DC simulations (empty circles) along with the values of *C*_*sat*_ as a function of temperature estimated through the TI-*C*_*sat*_ (red symbols) and TI-*CC* (blue symbols) schemes. The concentrations in the dilute phase evaluated using both TI approaches (and integrated from the closest temperature to the critical one, *T*_*c*_, depicted by cross symbols) remarkably match among themselves, and with the estimated values from DC simulations at moderately high temperatures. Importantly, we note that the calculation of *C*_*sat*_ fully relies on how accurately the coexisting densities have been calculated via DC simulations at the highest temperature (Fig. 1). While determining the saturation concentration via DC simulations at temperatures near *T*_*c*_ is straightforward, its evaluation at sub-physiological temperatures (*i*.*e*., below 310K, as commonly performed experimentally) is hardly attainable due to the extremely low coexisting concentrations of proteins in the dilute phase (usually below 0.1 mM). For instance, in FUS at 330 K, the average probability of finding a single protein in the corresponding volume of the dilute phase in a DC simulation is approximately one third (*i*.*e*., one every three recorded frames). If this probability is instead estimated from the *C*_*sat*_ value at 275 K, and considering the standard volume of a DC simulation, the likelihood of observing a single protein in the dilute phase decreases by a factor of *∼*10^*−*3^, corresponding to an expected occurrence every *∼*5 *µ*s on average. Therefore, to ensure reasonable sampling with at least 5 to 10 such events, the simulation timescale required for accurately estimating *C*_*sat*_ would need to span over 25 *µ*s. In Fig. S1 of the SM, we show how the obtained *C*_*sat*_’s as a function of temperature crucially depend on the initial concentrations employed as starting points (Fig. 1a). Nonetheless, if such equilibrium concentrations are accurately computed, their extrapolation to lower temperatures, and thus towards lower concentrations, can be safely applied. Importantly, for obtaining accurate calculations using the TI-*CC* method, we have computed Δ*U* within intervals of 20 K. For the TI-*C*_*sat*_ method, the intervals at which *U* was computed along each isochore was 10 K. Overall, our results in Fig. 2 demonstrate that both approaches reliably predict *C*_*sat*_ from phase boundaries at higher temperatures, supporting their validity and consistency. This enables the estimation of *C*_*sat*_ at lower temperatures, where direct calculations via DC simulations are not feasible.

**FIG. 2.**
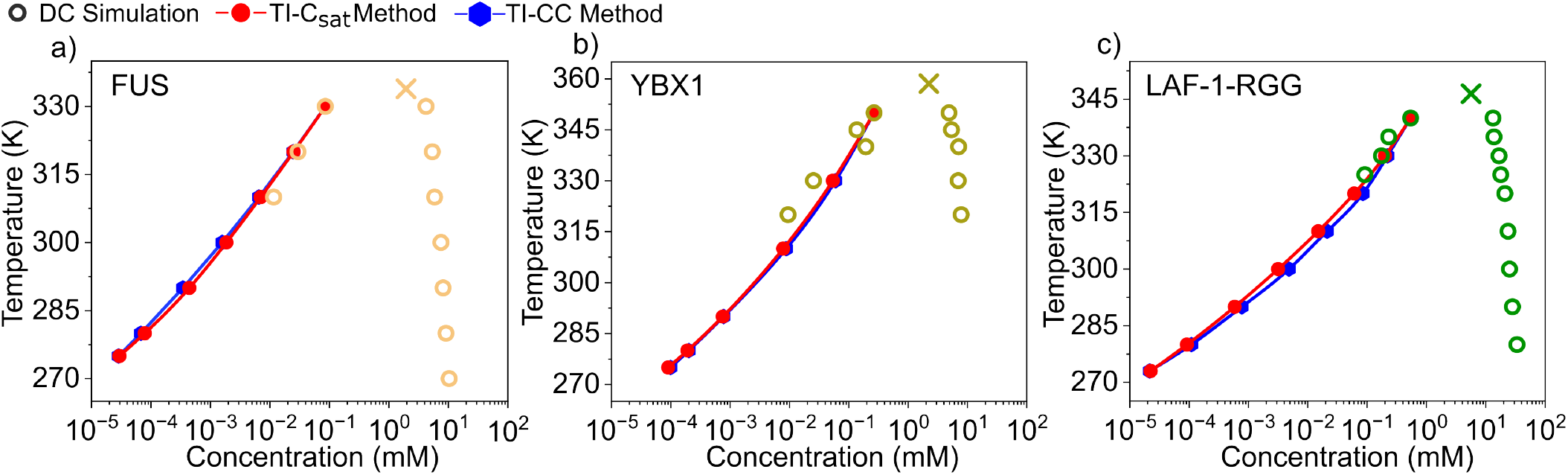
Phase diagrams of FUS (a), YBX1 (b), and LAF-1-RGG (c) in the temperature–concentration plane obtained from DC simulations (empty symbols) using the Mpipi-Recharged model. Cross symbols indicate the critical solution temperature evaluated through the law of rectilinear diameters and critical exponents^124^ using DC simulations. Red and blue filled symbols represent *C*_*sat*_ at different temperatures calculated via TI-*C*_*sat*_ and TI-*CC*, respectively, starting from the closest temperature to the critical point.

### B. Benchmark of Mpipi-Recharged phase diagrams vs. in vitro protein saturation concentrations

The biomolecular building blocks constituting condensates are commonly proteins with several sequence domains, including globular regions and IDRs, that enable the dynamic formation of intermolecular transient interactions which shape the stability and material properties of biomolecular condensates^125–127^. The protein sequence, in terms of amino acid composition, patterning, structure, and length crucially regulates the phase behaviour, explaining the high sensitivity of *C*_*sat*_ to subtle changes across the sequence, such as mutations and/or post-translational modifications^30,46,62,65,67,86,128^. To demonstrate the range of applicability of our proposed methods in combination with the Mpipi-Recharged for predicting biocondensate phase behaviour, we systematically compare *in vitro* phase diagrams and measurements of the saturation concentration^14,46,65,67,129^ with predictions of the Mpipi-Recharged combining DC simulations and the TI-*C*_*sat*_ approach.

Experimental studies have recently measured the saturation concentration (and even full phase diagrams) of multiple mutant variants of the LCD of hnRNPA1^65,67^— a multifunctional RNA-binding protein involved in various aspects of RNA metabolism, and linked to several neurodegenerative disorders^11^. Importantly, the specific mutations introduced across the sequence critically alter the phase separation propensity, providing a comprehensive analysis of the role of the amino acid nature, in terms of charge, polarity, hydrophobicity, or aromaticity in the ability to form condensates. These experimental phase diagrams and *C*_*sats*_ allow us to carry out a thorough comparison of the performance of the Mpipi-Recharged in describing protein phase behaviour. In Fig. 3, we present the phase diagrams obtained for 15 different hnRNPA1-LCD mutants showing in a circular diagram the sequence residue composition (in percentage) highlighting the most representative mutated residues across the set. This includes charged (lysine (K), arginine (R), aspartic acid (D), glutamic acid (E)), aromatic (tyrosine (Y), tryptophan (W), and phenylalanine (F)), and non-polar amino acids such as proline (P). Some other abundant amino acids such as serine (S) or glycine (G) have been excluded, along with those whose presence remains unchanged across the mutants, such as asparagine (N) or glutamine (Q) among others. In Fig. S3 of the SM we also provide a schematic depiction of the position of the aromatic, positively and negatively charged residues across the different variants.

**FIG. 3.**
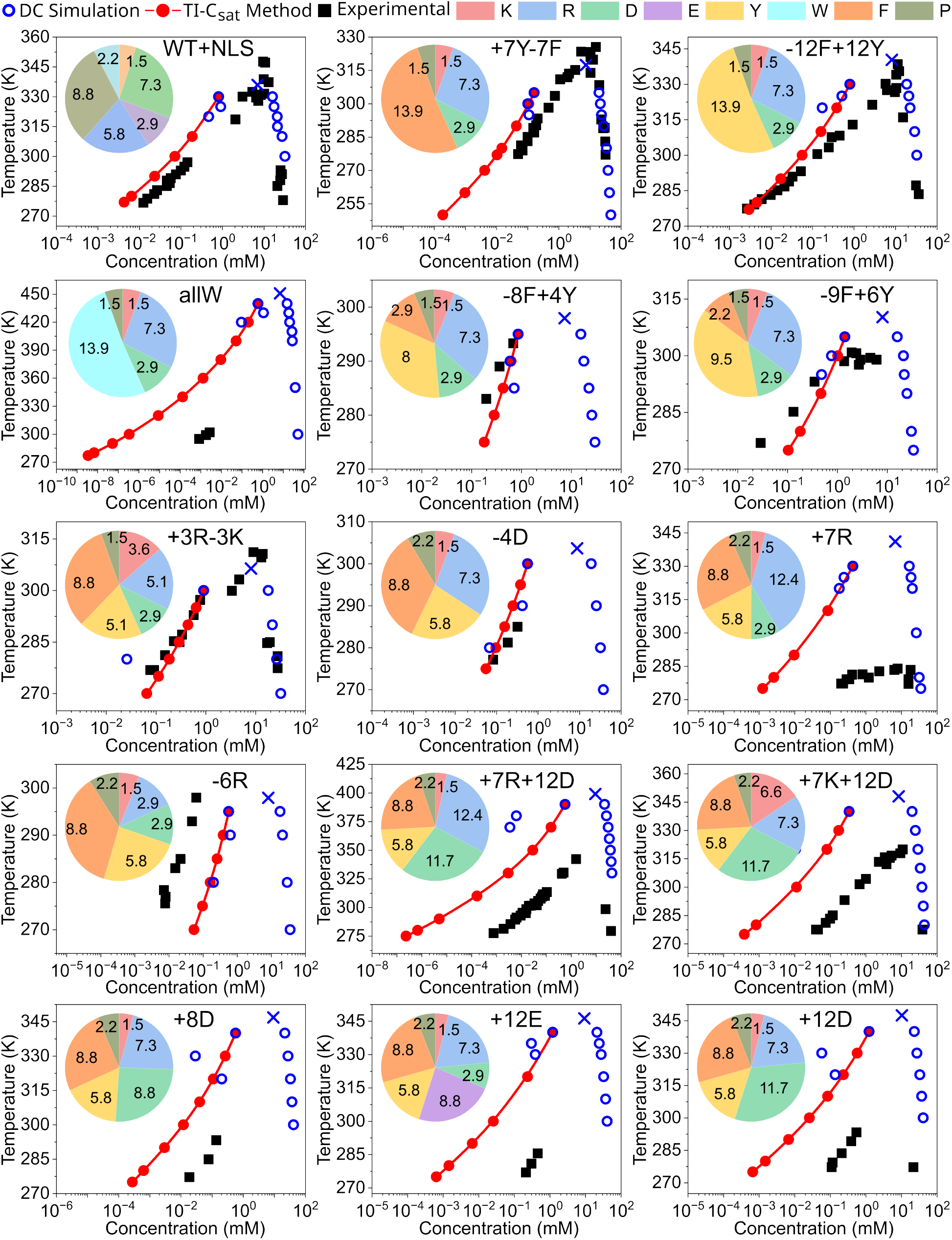
Phase diagrams in the temperature–concentration plane for different hnRNPA1-LCD mutants obtained from DC simulations (blue symbols), TI-*C*_*sat*_ calculations (red solid circles), and from *in vitro* experiments^65,67^ (black squares). Each panel includes a colour-coded circle diagram illustrating the amino acid composition (in percentage) of each corresponding sequence focusing on the most representative mutated residues across the set.

The phase diagrams predicted by the Mpipi-Recharged quantitatively match *in vitro* results for the WT+NLS sequence and the +7Y-7F, -12F+12Y, -8F+4Y, -9F+6Y, +3R-3K, and -4D mutants. For the -6R, +7R+12D, and +7K+12D variants it provides a reasonable estimation of the critical solution temperature but it underestimates *C*_*sat*_ for both +7R+12D, and +7K+12D, and moderately overestimates it for the -6R sequence. Finally, for the highly charged mutated variants (including +7R, +8D, +12E, or +12D) and allW the model underestimates the saturation concentration by *∼*2 orders of magnitude. The Mpipi-Recharged has consistently demonstrated to provide temperature–concentration phase diagrams that near-quantitatively agree with experimental *in vitro* measurements^68,69,108,123,130^. However, the uncertainty associated with the calculation of *C*_*sat*_ near the critical temperature via DC simulations—reaching up to half an order of magnitude for some cases^69^—makes challenging an exact comparison. Nevertheless, despite the associated uncertainty to the starting integration molar volume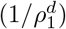, our TI routes enable the direct comparison at low temperatures, which for most of the sequences, would be unattainable through DC simulations. In that respect, Fig. 3 highlights this problem even at physiological temperatures, where the near-zero concentration of proteins in the dilute phase observed in DC simulations leads to *C*_*sat*_ values that can fluctuate by up to two orders of magnitude within just a few kelvins (*e*.*g*., the +12D mutant).

We identify the most pronounced discrepancies between our modelling predictions and *in vitro* results for the mutants with higher concentration of charged amino acids: +7R, +8D, +12E, and +12D. Due to the implicit treatment of ions and water, coarse-grained models of residue-resolution often struggle in describing the phase behaviour of highly charged sequences^69,108^. Nevertheless, the Mpipi-Recharged introduces a pair-specific asymmetric Yukawa electrostatic potential informed by atomistic simulations, to improve the description of highly charged systems. While previous coarse-grained models of similar resolution overestimate the critical solution temperature of charged protein-RNA condensates by hundreds of kelvin^47,112^, the Mpipi-Recharged shows a maximum offset in *T*_*c*_ of *∼*40K. Importantly, beyond the net charge of the protein, and the total number of charged residues, the distribution along the sequence (*i*.*e*., blockiness) plays a major role in the phase behaviour^45,46,65,67^. We have analysed the charge blockiness (see Section SVII of the SM) of the sequences with a highest concentration of charged residues. Nevertheless, as shown in Fig. S4, we cannot ascribe higher critical solution temperatures to mutant variants with higher degree of blockiness since the overall differences among the variants are modest. Instead, the mutation of lysines by arginines strongly enhances the ability to undergo phase-separation as shown for the +7R+12D and +7K+12D variants, and previously reported for different RNA-peptide complex coacervates^14,131^.

To further test the applicability of our methods—in particular we are using the TI-*C*_*sat*_ route, but as shown in Fig. 2 they both provide equivalent results—we now extend our analysis to different wild-type multi-domain proteins featuring both intrinsically disordered and globular domains. We combine DC simulations and TI-*C*_*sat*_ for calculating the saturation concentration of hnRNPA1, TDP-43, G3BP1, FUS, and YBX1. These RNA-and DNA-binding proteins are involved in the emergence of stress granules, and specifically, hnRNPA1, TDP-43, and FUS have been shown to promote a further liquid-to-solid transition into aberrant solid-like states linked to several neurodegenerative pathologies including ALS and FTD^6,11,136-140^. Therefore, a precise characterisation of their phase behaviour is fundamental to further advance our understanding on the onset these condensate-related pathologies. In Fig. 4a-e, we show the phase diagram from DC simulations (empty circles) along with the *C*_*sat*_ values from TI-*C*_*sat*_ (red solid circles) and the experimental saturation concentrations at physiological salt conditions (black triangles). We find a remarkable agreement between the predicted *C*_*sat*_ values and *in vitro* measurements^11,80,138^ across all systems, with the exception of TDP-43, where *C*_*sat*_ is overestimated by an order of magnitude. The highly specific *α*-*α*-helical interactions within the LCD of TDP-43 makes this sequence particularly difficult to be described. Nevertheless, in a previous study the Mpipi-Recharged has been shown to successfully describe its single-molecule radius of gyration, its reentrant phase behaviour in presence of poly-Uridine RNA and arginine-rich peptides, and its propensity to undergo condensate ageing over time^130^. Overall, these results from Fig. 4 demonstrate that: (1) the TI routes enable the direct comparison of experimental *vs*. modelling predictions by reducing the uncertainty in the calculation of the protein concentration in the dilute phase; and (2) the Mpipi-Recharged reproduces with significant accuracy the experimental saturation concentration of multi-domain proteins known to form condensates.

**FIG. 4.**
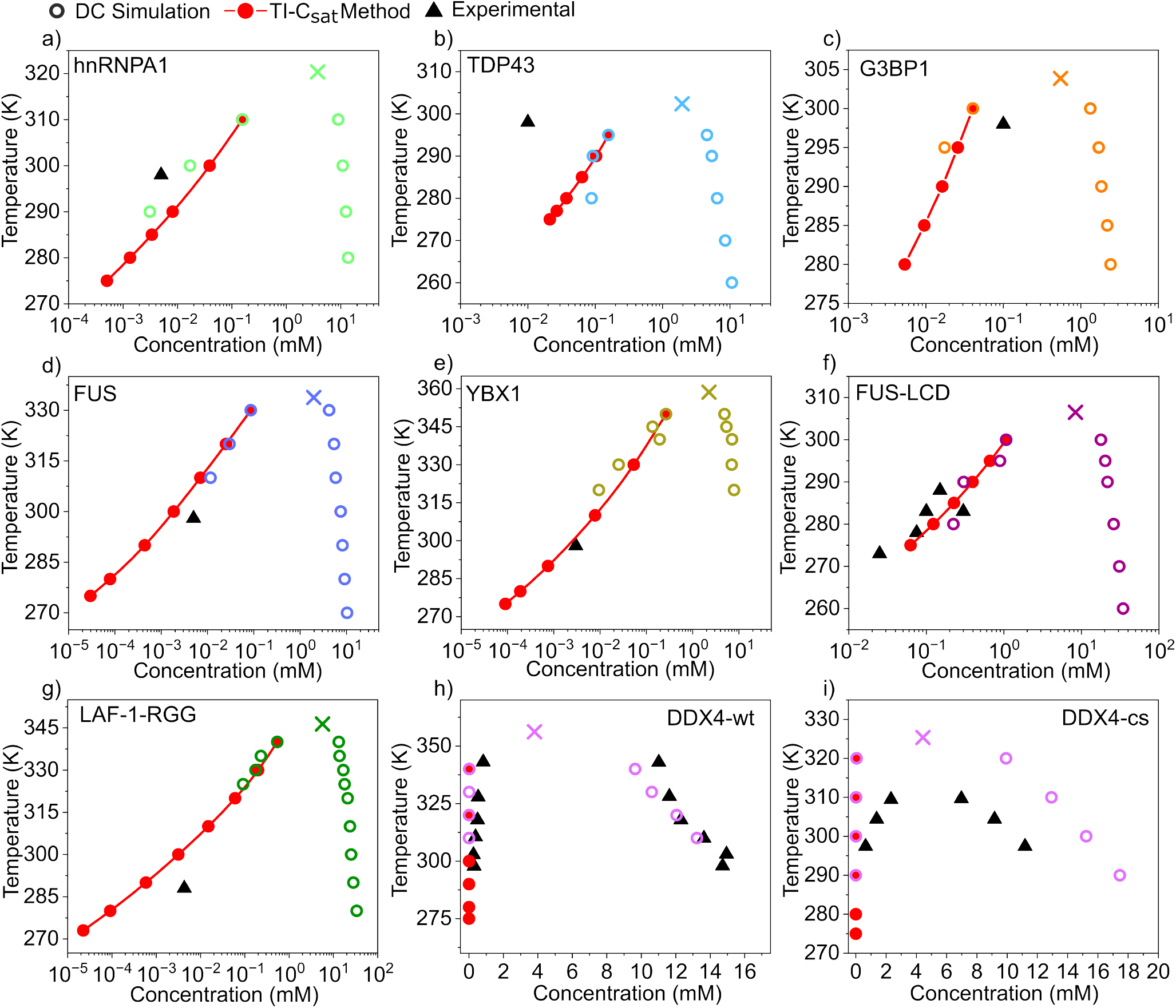
Phase diagrams in the temperature–concentration plane obtained through DC simulations of the Mpipi-Recharged (empty circles) and the TI-*C*_*sat*_ scheme (red solid circles) along with experimental saturation concentrations (black triangles) of various proteins including hnRNPA1^11^, TDP-43^48,49,80^, G3BP1^132^, FUS^49,80^, YBX1^133^, FUS-LCD^134^, LAF1-RGG^121^, and the wild-type (wt) and charge-scrambled (cs) sequences of DDX4^135^. Cross symbols depict the estimated critical solution temperature through the laws of rectilinear diameters and critical exponents using DC simulations.

Furthermore, we also apply the TI-*C*_*sat*_ scheme to IDPs including FUS-LCD, LAF-1-RGG, DDX4 wild-type (DDX4-wt), and the charged-scramble variant of DDX4 (DDX4-cs). The predicted phase diagrams along with the calculated *C*_*sat*_ values are shown in Fig. 4f-i. For FUS-LCD, we find an exceptional agreement between our simulations and *in vitro* saturation concentrations from previous works^127,134,141^. For LAF-1-RGG the thermodynamic integration method enables the direct comparison of *C*_*sat*_ at the experimental temperature (285 K)^121^, which would be unattainable through DC simulations at such temperature. The predicted value matches the experimental *C*_*sat*_ within half an order of magnitude, which approximately corresponds to the uncertainty of the method associated to the DC estimation at high temperature. Finally, the calculated coexistence lines for DDX4-wt and DDX4-cs are compared to experimental *in vitro* values^135^. Our results show good agreement between the simulated and experimental critical solution temperatures—especially for the DDX4 wild-type sequence (Fig. 4h)—with a maximum deviation of *∼*15K for DDX4-cs. Moreover, the concentration of both co-existing phases in both variants is fairly recovered by the Mpipi-Recharged, with particularly excellent predictions for the wild-type sequence. Nevertheless, a precise comparison between *C*_*sat*_ values as a function of temperature is hard to infer since the experimental data^135^ were not reported in *log* scale. Importantly, we note that both *in vitro* phase diagrams of DDX4 were carried out at a salt concentration different from the physiological one^14^ (100mM of NaCl). This increases the challenge for implicit-solvent CG models, as the Mpipi-Recharged, which have been initially parameterized for 150mM of NaCl despite the fact that they rescale their charge as a function of the ionic strength. Our results provided here show the significant improvement of the Mpipi-Recharged with respect to previous models in capturing the phase diagram of DDX4 and its charge-scramble variant^96,108,142^. In that respect, the overall charge of the protein, and more precisely the specific patterning of the charged residues in these two sequences, control the electrostatic interactions in a non-trivial manner which drastically modulates their phase diagram.

### C. C_sat_ predictions from explicit Mpipi-Recharged simulations and a machine-learning predictor

Molecular simulations have increasingly proven to be essential in understanding the microscopic processes involved in the formation of membraneless organelles through LLPS^70,76,106,108,143^. The continuous improvement of coarse-grained parameterizations to describe phase-separation^106,108,144,145^ has enabled successful predictions of multiple factors governing protein phase behaviour. In this respect, our TI methods offer novel opportunities to directly test modelling MD predictions against *in vitro* experimental measurements of a key parameter controlling LLPS: the saturation concentration. Nevertheless, alternative approaches which do not entail the performance of explicit MD simulations, such the use of machine-learning (ML) predictors, have been proven to be effective in providing accurate estimations of parameters determining LLPS such as the critical solution temperature, IDP conformational ensembles^146^, or saturation concentrations^71^. In this section, we compare explicit MD predictions of *C*_*sat*_ using the Mpipi-Recharged force field with those from an ML predictor^71^ based on the CALVADOS2 model^106^.

Since the CALVADOS2 ML-predictor^71^ only provides saturation concentration values at 293 K and 150 mM of NaCl, we first focus on the studied sequences for which values of *C*_*sat*_ are available under such conditions. These include all variants of hnRNPA1-LCD (Fig. 3) for which *C*_*sat*_ was reported at 293 K, and three additional variants, *i*.*e*. W^*−*^, FtoW, and YtoW from Ref.^67^ (see SM for further details on their sequences). As shown in Fig. 5a, both methods deliver reasonable estimates of *C*_*sat*_ for most of the variants. Strikingly, both models fail to accurately describe *C*_*sat*_ for the +12D sequence, highlighting the challenge of modelling highly charged sequences, where the interplay between electrostatic repulsion and attraction is particularly difficult to model at coarse-grained level^108^. Additionally, the Mpipi-Recharged significantly deviates in predicting *C*_*sat*_ of -6R, as evidenced in Fig. 3, and the CALVADOS2 ML-predictor in describing W^*−*^. We quantify the mean deviation of both computational datasets by calculating the quadratic average deviation (*D*) respect to the experimental reference values (please see Section SVIII in the SM for further details on this calculation). While both *D* values are relatively low (Fig. 5a)—reinforcing the realism of both approaches—the Mpipi-Recharged shows a slightly lower deviation respect to the experimental dataset inferred from Refs.^65,67^. We note that both methods— the ML-predictor^71^ and explicit MD simulations using DC and TI calculations—have several advantages and disadvantages. While the ML predictor is straightforward to use and computationally efficient—capable of estimating *C*_*sat*_ values for an entire proteome within a few hours—explicit MD simulations provide valuable molecular and mechanistic insights into the key intermolecular interactions driving LLPS, as well as into condensate architecture, surface free energy, and the conformational ensemble of proteins in both phases. However, these simulations require a significantly higher computational cost^68,69,147^. Hence, both approaches are fully complementary and contribute to elucidating across different scales—from an entire proteome to a particular protein segment interaction—the molecular factors governing condensate phase behaviour.

**FIG. 5.**
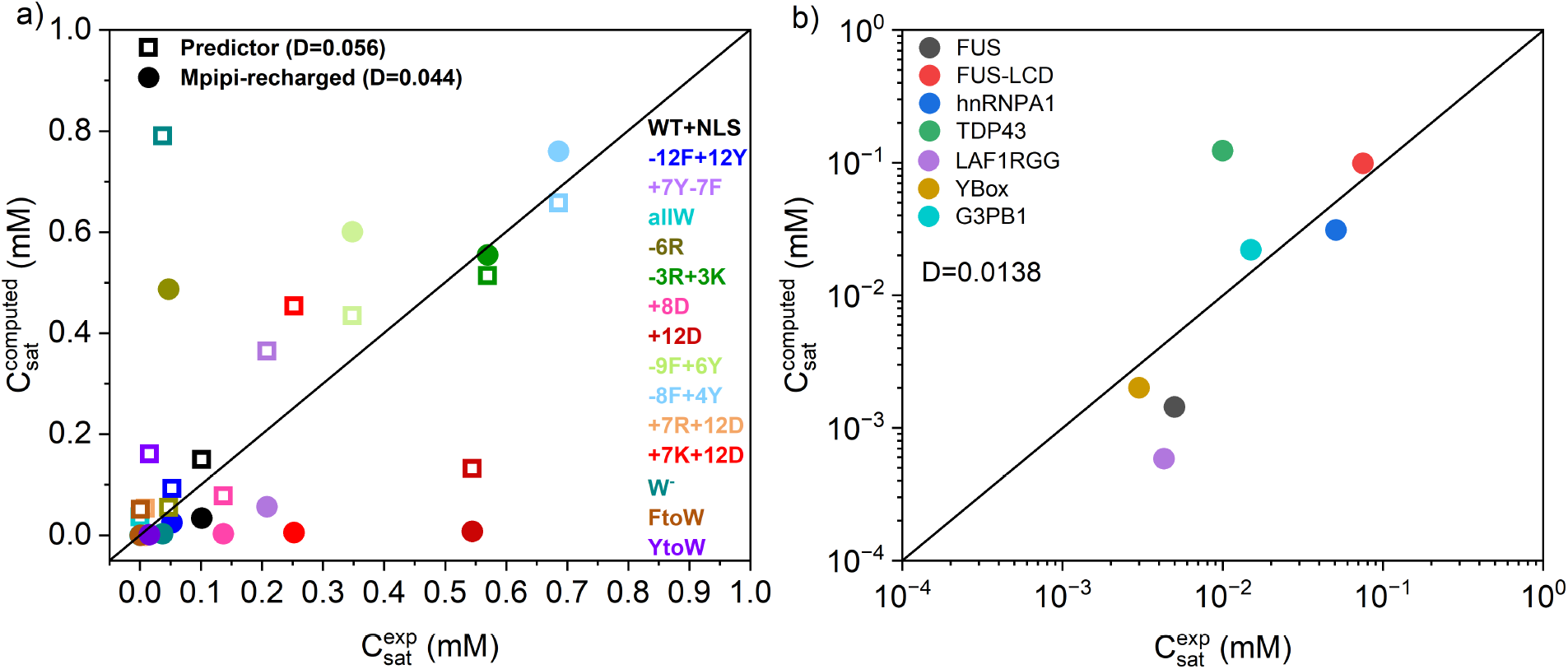
(a) Predicted saturation concentration 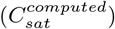 against *in vitro* 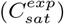 values for multiple mutated variants of hnRNPA1-LCD using the Mpipi-Recharged in combination with DC simulations and the TI-*C*_*sat*_ scheme (solid circles) and the CALVADOS2 ML-predictor^71^ (empty squares). The average quadratic deviation (*D*) of each modelling dataset from the experimental reference values^65,67^ is included in the legend. The black solid line represents a perfect match between experiments and simulations. (b) Comparison of the saturation concentration predicted by the Mpipi-Recharged against experimental protein saturation concentrations taken from Fig. 4 (black triangles).

Finally, in Fig. 5b we compare *C*_*sat*_ values obtained using the Mpipi-Recharged (and the TI-*C*_*sat*_ method) against experimental saturation concentrations of proteins that were not measured at 293K, and therefore cannot be evaluated through the CALVADOS2 ML-predictor^71^. Importantly, we note that while the experimental *C*_*sat*_ temperature is not constant among all these sequences (*i*.*e*., it ranges from 285 to 300K), the comparison between simulations and experiments has been performed at the same precise temperature for each case. Our results demonstrate a fair correlation between Mpipi-Recharged predictions and *in vitro* results^11,48,49,80,121,132–135,148^ in which only TDP-43 and LAF-1-RGG moderately deviate from the experimental value. Overall, by combining TI methods and DC simulations, we show a powerful framework to directly compare modelling and experimental saturation concentrations contributing to the continuous improvement and validation of biomolecular coarse-grained models.

## CONCLUSIONS

In this study, we introduce two computational methods for determining the saturation concentration of biomolecules undergoing phase separation. We first design a scheme (TI-*C*_*sat*_) in which a series of thermodynamic integration relations are applied and internal energies are computed as a function of temperature for bulk phases, enabling the calculation of arbitrarily low equilibrium saturation concentrations in coexistence with phase-separated condensates. Second, by using the Clausius-Clapeyron equation, we derive another approach (TI-*CC*) for evaluating *C*_*sat*_ at sub-physiological temperatures. This framework merely requires measuring internal energies in bulk phases at coexistence densities, initially determined from DC simulations, and subsequently obtained from the method itself. Importantly, despite the two methods being substantially different, both rely on the key assumption that proteins in the dilute phase behave like an ideal gas, exhibiting negligible intermolecular interactions. Both methods circumvent a major limitation of DC simulations in determining the concentration upon which phase-separation occurs in protein condensates^69,72,73^. The extremely low saturation concentrations of proteins to undergo LLPS— typically ranging from 1 to 200 *µ*M^12,127^—makes unfeasible its direct calculation through standard DC simulations, especially at moderately low temperatures (*i*.*e*., below 300 K), where most of the *in vitro* experimental measurements of *C*_*sat*_ are performed. Thus, our two methods—which provide equivalent predictions (Fig. 2)—enable the direct validation of *C*_*sat*_ modelling predictions *vs. in vitro* measurements (Figs. 3 and 4) in a straightforward and computationally feasible manner.

By using the TI-*C*_*sat*_ method, we have evaluated the protein saturation concentration dependence with temperature for over 20 different proteins, including purely intrinsically disordered proteins and multi-domain proteins—with both globular and disordered regions— which are known to undergo phase-separation. We have used the residue-resolution Mpipi-Recharged model recently developed by us^108^, which introduces a pair-specific asymmetric Yukawa electrostatic potential, informed by atomistic simulations, to compensate for the loss of explicit ions and water upon the coarse-graining. Having validated our thermodynamic integration frameworks for different protein sequences (Fig. 2), we have compared the predicted full phase diagram by the Mpipi-Recharged for 15 different mutants of the low-complexity domain of hnRNPA1 with *in vitro* predictions of either the phase diagram, or the protein saturation concentration as a function of temperature (Fig. 3). The model remarkably predicts the critical solution temperature and *C*_*sat*_ for most of the hnRNPA1-LCD variants, except for those with a significantly high concentration of mutated charged amino acids. Nevertheless, even for those sequences, the maximum offset in the critical solution temperature does not exceed *∼*40 K, while *C*_*sat*_ does not differ in more than two orders of magnitude from *in vitro* measurements^65,67^. Because protein saturation concentrations can vary by more than five orders of magnitude depending on factors like specific sequence mutations, temperature, pH, and salt concentration, predictions that fall within the correct order of magnitude are already highly informative. In that sense, our thermodynamic integration methods improve the accuracy of these calculations, particularly at low temperatures, where sufficient sampling through DC calculations is generally unattainable.

Furthermore, we have tested our predictions for several IDPs and multi-domain RNA-and DNA-binding proteins involved in the formation of stress granules, whose liquid-to-solid transition upon phase-separation has been linked to ALS and FTD diseases^6^ (Fig. 4). We find a remarkable agreement between the predicted *C*_*sat*_ and *in vitro* measurements across all systems^11,49,80,121,132-135^, with the exception of TDP-43^48,80^, in which highly specific *α*-*α*-helical interactions within its LCD makes this sequence particularly challenging to be modelled. Nevertheless, even for TDP-43 the offset in *C*_*sat*_ with the experimental value is below an order of magnitude. These results highlight that: (1) our TI methods enable direct comparison between experimental and modelling predictions in a temperature and concentration range that is not accessible through DC simulations; and (2) the Mpipi-Recharged reproduces the experimental saturation concentration of numerous condensate-forming multi-domain proteins with significant accuracy.

Finally, we have benchmarked our explicit MD simulations using the Mpipi-Recharged with a machine-learning predictor based on the successful residue-resolution CAL-VADOS2 coarse-grained model^71^ (Fig. 5). We show that both approaches yield reasonable estimates of *C*_*sat*_ for most hnRNPA1-LCD mutant variants with available experimental data under the conditions for which the ML predictor was trained (293 K and physiological salt conditions). Interestingly, both models have difficulty capturing *C*_*sat*_ for highly charged engineered variants, such as the +12D sequence, highlighting the challenge of accurately modelling the interplay between electrostatic repulsion and attraction, which has been recently shown to improve through the implementation of asymmetric potentials that soften electrostatic repulsion^108^. By quantifying the mean deviation of both computational datasets with respect to the experimental reference values, we find that both approaches provide realistic predictions of *C*_*sat*_ values with a slightly lower deviation of the Mpipi-Recharged for the tested dataset. Importantly, we remark that both methods—the ML-predictor^71^ and explicit MD simulations using DC and TI calculations—have several advantages and drawbacks, such as ML-predictors being straightforward to use and highly computationally efficient, but incapable to generate by construction mechanistic and molecular insights of the key sequence signatures driving condensation, and the DC/TI schemes yielding these insights but at a higher computational cost. In summary, our TI methods presented here represent a powerful tool to compare modelling and experimental predictions, contributing to the improvement and validation of biomolecular coarse-grained models to investigate the physicochemical mechanisms regulating condensate formation, and enhancing the scope of computer simulations in this field. Therefore, these approaches not only complement existing experimental techniques, but also provide a robust framework for simulating and predicting LLPS in complex protein systems.

## ACKNOWLEDGEMENTS

E.P. acknowledges funding from an FPI scholarship associated to the Spanish scientific plan and committee for research: PID2022-136919NA-C33. R.C.G, and A.T. acknowledge funding from the UK Research and Innovation (UKRI) Engineering and Physical Sciences Research Council (EPSRC) under the UK Government’s guarantee scheme (EP/Z002028/1), following successful evaluation by the ERC (Consolidator Grant awarded to R.C.G.) under the European Union’s Horizon Europe research and innovation programme. A. F. acknowledges funding from the Ramon y Cajal fellowship (RYC2021-030937-I) and Spanish National Grant (PID2022-136919NA-C33). J. R. E. acknowledges funding from Emmanuel College, the University of Cambridge, and the Ramon y Cajal fellowship (RYC2021-030937-I). J. R. E. and F. G. also acknowledge the Spanish scientific plan and committee for research; project reference PID2022-136919NA-C33. E. S. acknowledges project PID2022-136919NB-C31 from the the Spanish MCIU. This work has been performed using resources provided by the Cambridge Tier-2 system operated by the University of Cambridge Research Computing Service (http://www.hpc.cam.ac.uk) funded by EPSRC Tier-2 capital grant EP/P020259/1-CS170. This work has also been performed using resources provided by Archer2 (https://www.archer2.ac.uk/) funded by EPSRC Tier-2 capital grant EP/P020259/e829. The authors also thankfully acknowledge RES computational resources provided by Mare Nostrum 5 through the activities 2024-3-0001 and 2025-1-0009.

### APPENDIX FORMULATION OF TI-C_SAT_ METHOD

The temperature–density phase diagram establishes the limiting boundaries for observing phase separation, and it sets the range of temperatures for building our thermodynamic route. Starting from the equilibrium densities of the binodal at temperature *T*_1_, we set a target lower temperature *T*_2_ (Fig. 1a). The thermodynamic connection between both states is achieved through an isochoric process 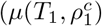 to 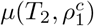 and 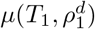 to 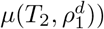 followed by an isothermal transformation 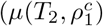 to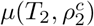; and 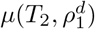 to 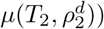 that intersects the binodal. At any point in the coexistence line for a given temperature, the chemical potential is the same for both phases since the system is at equilibrium, *i*.*e*.:

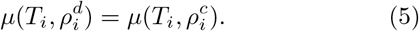

The above relation can be particularised for the initial state at *T*_1_ and the target state at *T*_2_. Thus, we can calculate the difference as:

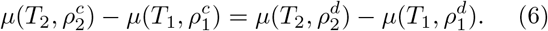

From the previous equation, we can incorporate the intermediate steps by first considering the isochoric process, followed by the isothermal process. Therefore, we add the terms -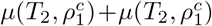 and -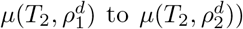 to the left hand and right hand sides of Eq. 6, respectively, to obtain:

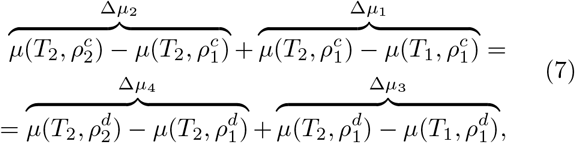

and thus:

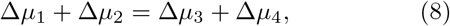

where Δ*µ*_1_ and Δ*µ*_3_ represent the variation in the chemical potential along the isochores for the condensed and dilute phases, and Δ*µ*_2_ and Δ*µ*_4_ correspond to the difference in chemical potential along the isotherms for the condensed and dilute phases, respectively (see Fig. 1b).

Since we aim to determine 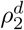, we can relate this density with Δ*µ*_4_, which represents an isothermal expansion of the dilute phase. Assuming ideal gas behaviour for the dilute phase:

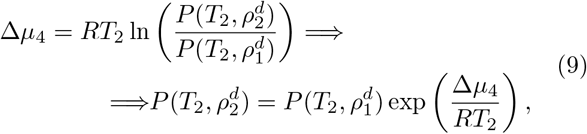

and knowing that for an ideal gas 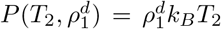 and 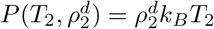, from Eq. 9 we obtain:

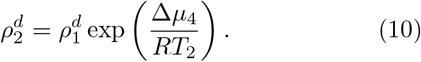

In this way, from Eq. 10 we can obtain 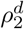 (our target) if we first determine Δ*µ*_4_ as:

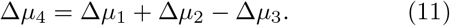

We can determine the isothermal compression of the condensed phase represented by Δ*µ*_2_ using Eq. 12. However, the pressure from 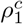 to 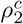 barely changes across phase-separated condensate simulations, and even in some cases negatives pressures can be obtained unless the sampling is large. Thus, we can assume that Δ*µ*_2_ *≈* 0:

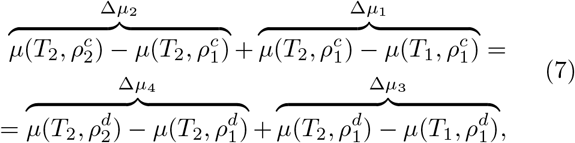

Therefore Δ*µ*_4_ can be calculated as:

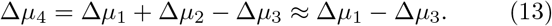

The chemical potential difference Δ*µ*_1_ and Δ*µ*_3_ can be obtained using the general expression:

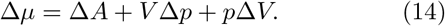

Due to the constant volume along the isochoric path and knowing that the pressure change is negligible, Δ*µ*_1,3_ can be approximated to Δ*A*_1,3_, *i*.*e*.:

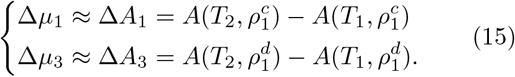

On the other hand, 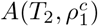 and 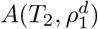 can be straightforwardly obtained from the thermodynamic integration relations^117,149^:

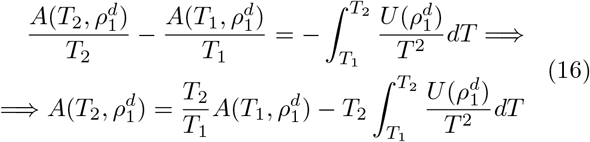

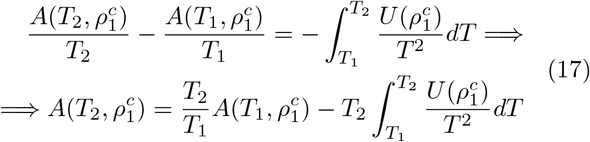

By substituting Eqs. 16 and 17 in Eq. 15, we obtain:

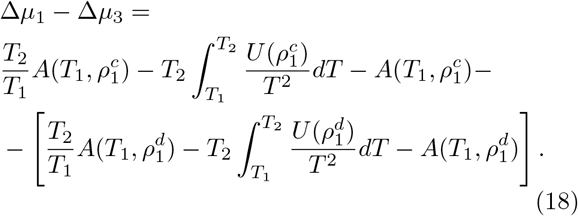

Hence, since 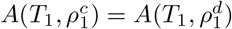 given that the equilibrium pressure is close to zero:

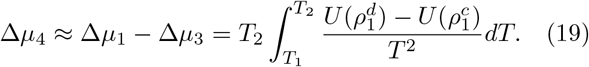

Now, replacing Δ*µ*_4_ in Eq. 10 we obtain the relation:

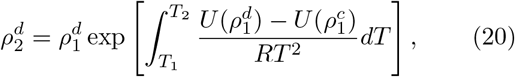

where 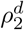 is the coexistence density at a temperature *T*_2_ lower than *T*_1_ that we aim to determine, 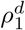 is the coexistence density for the dilute phase determined at *T*_1_, *R* is the ideal gas constant, *T* is the temperature and 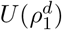 and 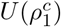 correspond to the internal energy at each temperature across the thermodynamic integration pathway (Fig. 1b).

## REFERENCES

1 S. F. Banani, H. O. Lee, A. A. Hyman, and M. K. Rosen, Nature Reviews Molecular Cell Biology 18, 285 (2017).

2 S. Alberti, Current Biology 27, R1097 (2017).

3 A. A. Hyman and K. Simons, Science 337, 1047 (2012).

4 A. A. Hyman, C. A. Weber, and F. Jülicher, Annual Review of Cell and Developmental Biology 30, 39 (2014).

5 A. Patel, H. O. Lee, L. Jawerth, S. Maharana, M. Jahnel, M. Y. Hein, S. Stoynov, J. Mahamid, S. Saha, T. M. Franzmann, et al., Cell 162, 1066 (2015).

6 B. Portz, B. L. Lee, and J. Shorter, Trends in Biochemical Sciences 46, 550 (2021).

7 B. Wang, L. Zhang, T. Dai, Z. Qin, H. Lu, L. Zhang, and F. Zhou, Signal Transduction and Targeted Therapy 6 (2021), 10.1038/s41392-021-00678-1.

8 Y. Shin and C. P. Brangwynne, Science 357 (2017).

9 A. R. Strom, A. V. Emelyanov, M. Mir, D. V. Fyodorov, X. Darzacq, and G. H. Karpen, Nature 547, 241 (2017).

10 P. Li, S. Banjade, H.-C. Cheng, S. Kim, B. Chen, L. Guo, M. Llaguno, J. V. Hollingsworth, D. S. King, S. F. Banani, P. S. Russo, Q.-X. Jiang, B. T. Nixon, and M. K. Rosen, Nature 483, 336 (2012).

11 A. Molliex, J. Temirov, J. Lee, M. Coughlin, A. P. Kanagaraj, H. J. Kim, T. Mittag, and J. P. Taylor, Cell 163, 123 (2015).

12 S. Boeynaems, S. Alberti, N. L. Fawzi, T. Mittag, M. Polymenidou, F. Rousseau, J. Schymkowitz, J. Shorter, B. Wolozin, L. Van Den Bosch, et al., Trends in cell biology 28, 420 (2018).

13 S. F. Banani, A. M. Rice, W. B. Peeples, Y. Lin, S. Jain, R. Parker, and M. K. Rosen, Cell 166, 651 (2016).

14 J. P. Brady, P. J. Farber, A. Sekhar, Y.-H. Lin, R. Huang, A. Bah, T. J. Nott, H. S. Chan, A. J. Baldwin, J. D. Forman-Kay, et al., Proceedings of the National Academy of Sciences 114, E8194 (2017).

15 F. Frottin, F. Schueder, S. Tiwary, R. Gupta, R. Korner, T. Schlichthaerle, J. Cox, R. Jungmann, F. Hartl, and M. Hipp, Science 365, 342 (2019).

16 J. A. Riback, M. A. Bowman, A. M. Zmyslowski, C. R. Knoverek, J. M. Jumper, J. R. Hinshaw, E. B. Kaye, K. F. Freed, P. L. Clark, and T. R. Sosnick, Science 358, 238 (2017).

17 A. H. Fox, S. Nakagawa, T. Hirose, and C. S. Bond, Trends in Biochemical Sciences 43, 124 (2018).

18 S. Yang, S. T. Warraich, G. A. Nicholson, and I. P. Blair, The International Journal of Biochemistry & Cell Biology 42, 1408 (2010).

19 L. Yang, J. Gal, J. Chen, and H. Zhu, Proceedings of the National Academy of Sciences 111, 17809 (2014).

20 S. Keyport Kik, D. Christopher, H. Glauninger, C. W. Hickernell, J. A. M. Bard, K. M. Lin, A. H. Squires, M. Ford, T. R. Sosnick, and D. A. Drummond, Nature Communications 15, 3127 (2024).

21 F. Niss, L. Piñero-Paez, W. Zaidi, E. Hallberg, and A.-L. Ström, Molecular Neurobiology 59, 5236 (2022).

22 J. Gal, J. Chen, D.-Y. Na, L. Tichacek, K. R. Barnett, and H. Zhu, Molecular and Cellular Biology 39, e00052 (2019).

23 B. Al-Sady, H. Madhani, and G. Narlikar, Molecular Cell 51, 80 (2013).

24 L. F. S. Bonet, J. P. Loureiro, G. R. C. Pereira, A. N. R. Da Silva, and J. F. De Mesquita, PLOS ONE 16, e0258061 (2021).

25 M. Vendruscolo and M. Fuxreiter, Journal of Molecular Biology 434, 167201 (2022).

26 C. Vance, B. Rogelj, T. Hortobágyi, K. J. De Vos, A. L. Nishimura, J. Sreedharan, X. Hu, B. Smith, D. Ruddy, P. Wright, et al., Science 323, 1208 (2009).

27 M. Kamelgarn, J. Chen, L. Kuang, H. Jin, E. J. Kasarskis, and H. Zhu, Proceedings of the National Academy of Sciences 115, E11904 (2018).

28 S. Alberti and A. A. Hyman, Nature Reviews Molecular Cell Biology 22, 196 (2021).

29 S. Wegmann, B. Eftekharzadeh, K. Tepper, K. M. Zoltowska, R. E. Bennett, S. Dujardin, P. R. Laskowski, D. MacKenzie, T. Kamath, C. Commins, et al., The EMBO Journal 37, e98049 (2018).

30 S. Maharana, J. Wang, D. K. Papadopoulos, D. Richter, A. Pozniakovsky, I. Poser, M. Bickle, S. Rizk, J. Guillen-Boixet, T. M. Franzmann, M. Jahnel, L. Marrone, Y.-T. Chang, J. Sterneckert, P. Tomancak, A. A. Hyman, and S. Alberti, Science 360, 918 (2018).

31 N. L. Gotor, A. Armaos, G. Calloni, M. Torrent Burgas, R. M. Vabulas, N. S. De Groot, and G. G. Tartaglia, Nucleic Acids Research 48, 9491 (2020).

32 S. Torrino, W. M. Oldham, A. R. Tejedor, I. S. Burgos, L. Nasr, N. Rachedi, K. Fraissard, C. Chauvet, C. Sbai, B. P. O’Hara, et al., Cell 188, 447 (2025).

33 S. Qamar, G. Wang, S. J. Randle, F. S. Ruggeri, J. A. Varela, J. Q. Lin, E. C. Phillips, A. Miyashita, D. Williams, F. Ströhl, et al., Cell 173, 720 (2018).

34 X. Gui, F. Luo, Y. Li, H. Zhou, Z. Qin, Z. Liu, J. Gu, M. Xie, K. Zhao, B. Dai, et al., Nature Communications 10, 1 (2019).

35 Y. Sun, K. Zhao, W. Xia, G. Feng, J. Gu, Y. Ma, X. Gui, X. Zhang, Y. Fang, B. Sun, et al., Nature Communications 11, 1 (2020).

36 P. S. Tsoi, M. D. Quan, K.-J. Choi, K. M. Dao, J. C. Ferreon, and A. C. M. Ferreon, Protein Science 30, 1408 (2021).

37 S. Elbaum-Garfinkle, Y. Kim, K. Szczepaniak, C. C.-H. Chen, C. R. Eckmann, S. Myong, and C. P. Brangwynne, Proceedings of the National Academy of Sciences 112, 7189 (2015).

38 X.-M. Liu, L. Ma, and R. Schekman, elife 10, e71982 (2021).

39 L. Lim, Y. Wei, Y. Lu, and J. Song, PLOS Biology 14, 1 (2016).

40 P. Mohanty, A. Rizuan, Y. C. Kim, N. L. Fawzi, and J. Mittal, Protein Science 33, e4891 (2024).

41 T. Afroz, E.-M. Hock, P. Ernst, C. Foglieni, M. Jambeau, L. A. Gilhespy, F. Laferriere, Z. Maniecka, A. Plückthun, P. Mittl, et al., Nature Communications 8, 1 (2017).

42 A. G. Larson, D. Elnatan, M. M. Keenen, M. J. Trnka, J. B. Johnston, A. L. Burlingame, D. A. Agard, S. Redding, and G. J. Narlikar, Nature 547, 236 (2017).

43 M. M. Keenen, D. Brown, L. D. Brennan, R. Renger, H. Khoo, C. R. Carlson, B. Huang, S. W. Grill, G. J. Narlikar, and S. Redding, elife 10, e64563 (2021).

44 X. Jin, M. Zhou, S. Chen, D. Li, X. Cao, and B. Liu, Cellular and Molecular Life Sciences 79, 380 (2022).

45 N. Galvanetto, M. T. Ivanović, A. Chowdhury, A. Sottini, M. F. Nüesch, D. Nettels, R. B. Best, and B. Schuler, Nature 619, 876 (2023).

46 H. Yamazaki, M. Takagi, H. Kosako, T. Hirano, and S. H. Yoshimura, Nature Cell Biology 24, 625 (2022).

47 A. R. Tejedor, A. Garaizar, J. Ramírez, and J. R. Espinosa, Biophysical Journal 120, 5169 (2021).

48 L. McGurk, E. Gomes, L. Guo, J. Mojsilovic-Petrovic, V. Tran, R. G. Kalb, J. Shorter, and N. M. Bonini, Molecular Cell 71, 703 (2018).

49 W. M. Babinchak, R. Haider, B. K. Dumm, P. Sarkar, K. Surewicz, J.-K. Choi, and W. K. Surewicz, Journal of Biological Chemistry 294, 6306 (2019).

50 A. Wang, A. E. Conicella, H. B. Schmidt, E. W. Martin, S. N. Rhoads, A. N. Reeb, A. Nourse, D. Ramirez Montero, V. H. Ryan, R. Rohatgi, et al., The EMBO journal 37, e97452 (2018).

51 N. Farahi, T. Lazar, S. J. Wodak, P. Tompa, and R. Pancsa, International Journal of Molecular Sciences 22 (2021), 10.3390/ijms22063017.

52 K. Burke, A. Janke, C. Rhine, and N. Fawzi, Molecular Cell 60, 231 (2015).

53 R. Kobayashi and H. Nabika, Soft Matter 20, 5331 (2024).

54 M. Brzezinski, P. G. Argudo, T. Scheidt, M. Yu, E. Hosseini, A. Kaltbeitzel, E. A. Lemke, J. J. Michels, and S. H. Parekh, Biomacromolecules 26, 2060 (2025).

55 A. A. M. André and E. Spruijt, International Journal of Molecular Sciences 21 (2020), 10.3390/ijms21165908.

56 C. K. Patel, S. Singh, B. Saini, and T. K. Mukherjee, The Journal of Physical Chemistry Letters 13, 3636 (2022).

57 X. Zhang, H. Li, Y. Ma, D. Zhong, and S. Hou, APL Bioengineering 7, 021502 (2023).

58 H. Zhang, S. Shao, and Y. Sun, Biophysics Reports 8, 2 (2022).

59 S. Choi, S. Y. Chun, K. Kwak, and M. Cho, Physical Chemistry Chemical Physics 25, 9051 (2023).

60 M. Brzezinski, P. Argudo, J. Michels, and S. H. Parekh, Biophysical Journal 123, 445a (2024).

61 M. Loidolt-Krüger, Methods in Microscopy (2025).

62 S. Ray, N. Singh, R. Kumar, K. Patel, S. Pandey, D. Datta, J. Mahato, R. Panigrahi, A. Navalkar, S. Mehra, et al., Nature chemistry 12, 705 (2020).

63 M.-T. Wei, S. Elbaum-Garfinkle, A. S. Holehouse, C. C.-H. Chen, M. Feric, C. B. Arnold, R. D. Priestley, R. V. Pappu, and C. P. Brangwynne, Nature Chemistry 9, 1118 (2017).

64 F. Muzzopappa, J. Hummert, M. Anfossi, S. A. Tashev, D.-P. Herten, and F. Erdel, Nature Communications 13, 7787 (2022).

65 A. Bremer, M. Farag, W. M. Borcherds, I. Peran, E. W. Martin, R. V. Pappu, and T. Mittag, Nature Chemistry 14, 196 (2022).

66 I. Alshareedah, M. M. Moosa, M. Pham, D. A. Potoyan, and P. R. Banerjee, Nature Communications 12, 1 (2021).

67 I. Alshareedah, W. M. Borcherds, S. R. Cohen, A. Singh, A. E. Posey, M. Farag, A. Bremer, G. W. Strout, D. T. Tomares, R. V. Pappu, T. Mittag, and P. R. Banerjee, Nature Physics 20, 1482 (2024).

68 A. Feito, I. Sanchez-Burgos, A. Rey, R. Collepardo-Guevara, J. R. Espinosa, and A. R. Tejedor, Molecular Physics 122, e2425757 (2024).

69 A. Feito, I. Sanchez-Burgos, I. Tejero, E. Sanz, A. Rey, R. Collepardo-Guevara, A. R. Tejedor, and J. R. Espinosa, PLOS Computational Biology 21, e1012737 (2025).

70 G. Tesei, T. K. Schulze, R. Crehuet, and K. Lindorff-Larsen, Proceedings of the National Academy of Sciences 118, e2111696118 (2021).

71 S. von Bülow, G. Tesei, F. K. Zaidi, T. Mittag, and K. Lindorff-Larsen, Proceedings of the National Academy of Sciences 122, e2417920122 (2025).

72 M. J. Maristany, A. A. Gonzalez, J. R. Espinosa, J. Huertas, R. Collepardo-Guevara, and J. A. Joseph, eLife 13, RP99068 (2025).

73 F. Cao, S. von Bülow, G. Tesei, and K. Lindorff-Larsen, “A coarse-grained model for disordered and multi-domain proteins,” (2024).

74 G. L. Dignon, W. Zheng, R. B. Best, Y. C. Kim, and J. Mittal, Proceedings of the National Academy of Sciences 115, 9929 (2018).

75 M. Orozco, A. Pérez, A. Noy, and F. J. Luque, Chemical Society Reviews 32, 350 (2003).

76 B. Szala-Mendyk, T. M. Phan, P. Mohanty, and J. Mittal, Current Opinion in Chemical Biology 75, 102333 (2023).

77 G. L. Dignon, W. Zheng, Y. C. Kim, R. B. Best, and J. Mittal, PLoS Computational Biology 14, e1005941 (2018).

78 J.-M. Choi, A. S. Holehouse, and R. V. Pappu, Annual Review of Biophysics 49, 107 (2020).

79 A. Garaizar, J. R. Espinosa, J. A. Joseph, G. Krainer, Y. Shen, T. P. Knowles, and R. Collepardo-Guevara, Proceedings of the National Academy of Sciences 119, e2119800119 (2022).

80 G. Krainer, T. J. Welsh, J. A. Joseph, J. R. Espinosa, S. Wittmann, E. de Csillery, A. Sridhar, Z. Toprakcioglu, G. Gudiskyte, M. A. Czekalska, et al., Nature Communications 12, 1 (2021).

81 S. Blazquez, I. Sanchez-Burgos, J. Ramirez, T. Higginbotham, M. M. Conde, R. Collepardo-Guevara, A. R. Tejedor, and J. R. Espinosa, Advanced Science 10, 2207742 (2023).

82 S. Shi, L. Zhao, and Z.-Y. Lu, The Journal of Physical Chemistry Letters 15, 7280 (2024).

83 J. R. Espinosa, J. A. Joseph, I. Sanchez-Burgos, A. Garaizar, D. Frenkel, and R. Collepardo-Guevara, Proceedings of the National Academy of Sciences 117, 13238 (2020).

84 J. R. Espinosa, A. Garaizar, C. Vega, D. Frenkel, and R. Collepardo-Guevara, The Journal of Chemical Physics 150, 224510 (2019).

85 A. Garaizar, T. Higginbotham, I. Sanchez-Burgos, A. R. Tejedor, E. Sanz, and J. R. Espinosa, The Journal of Chemical Physics 157 (2022).

86 A. Garaizar, I. Sanchez-Burgos, R. Collepardo-Guevara, and J. R. Espinosa, Molecules 25 (2020), 10.3390/molecules25204705.

87 L. G. MacDowell and F. J. Blas, The Journal of Chemical Physics 131, 074705 (2009).

88 J.-M. Choi, F. Dar, and R. V. Pappu, PLOS Computational Biology 15, e1007028 (2019).

89 S. Das, A. Eisen, Y.-H. Lin, and H. S. Chan, The Journal of Physical Chemistry B 122, 5418 (2018).

90 T. S. Harmon, A. S. Holehouse, M. K. Rosen, and R. V. Pappu, elife 6, e30294 (2017).

91 S. Das, Y.-H. Lin, R. M. Vernon, J. D. Forman-Kay, and H. S. Chan, Proceedings of the National Academy of Sciences 117, 28795 (2020).

92 A. K. Lancaster, A. Nutter-Upham, S. Lindquist, and O. D. King, Bioinformatics 30, 2501 (2014).

93 M. Pu, B. Fu, and Y. J. Zhang, bioRxiv (2022), 10.1101/2022.07.25.501404.

94 Q. Liang, N. Peng, Y. Xie, N. Kumar, W. Gao, and Y. Miao, The EMBO Journal 43, 1898 (2024).

95 M. Monti, J. Fiorentino, D. Miltiadis-Vrachnos, G. Bini, T. Cotrufo, N. S. de Groot, A. Armaos, and G. G. Tartaglia, bioRxiv (2024), 10.1101/2024.07.19.602785.

96 R. M. Vernon, P. A. Chong, B. Tsang, T. H. Kim, A. Bah, P. Farber, H. Lin, and J. D. Forman-Kay, elife 7, e31486 (2018).

97 K. Y. Chin, S. Ishida, Y. Sasaki, and K. Terayama, BMC Bioinformatics 25, 143 (2024).

98 K. L. Saar, A. S. Morgunov, R. Qi, W. E. Arter, G. Krainer, A. A. Lee, and T. P. J. Knowles, Proceedings of the National Academy of Sciences 118, e2019053118 (2021).

99 P. Mullick and A. Trovato, Biomolecules 12 (2022), 10.3390/biom12121771.

100 S. Boeynaems, A. S. Holehouse, V. Weinhardt, D. Kovacs, J. Van Lindt, C. Larabell, L. Van Den Bosch, R. Das, P. S. Tompa, R. V. Pappu, et al., Proceedings of the National Academy of Sciences 116, 7889 (2019).

101 J. Sun, J. Qu, C. Zhao, X. Zhang, X. Liu, J. Wang, C. Wei, X. Liu, M. Wang, P. Zeng, X. Tang, X. Ling, L. Qing, S. Jiang, J. Chen, T. S. R. Chen, Y. Kuang, J. Gao, X. Zeng, D. Huang, Y. Yuan, L. Fan, H. Yu, and J. Ding, Nature Communications 15, 2662 (2024).

102 E. R. Kuechler, A. Huang, J. M. Bui, T. Mayor, and J. Gsponer, Biomolecules 13 (2023), 10.3390/biom13030527.

103 G. M. Ginell, R. J. Emenecker, J. M. Lotthammer, E. T. Usher, and A. S. Holehouse, bioRxiv (2024), 10.1101/2024.06.03.597104.

104 L. M. Blaabjerg, M. M. Kassem, L. L. Good, N. Jonsson, M. Cagiada, K. E. Johansson, W. Boomsma, A. Stein, and K. Lindorff-Larsen, Elife 12, e82593 (2023).

105 G. van Mierlo, J. R. Jansen, J. Wang, I. Poser, S. J. van Heeringen, and M. Vermeulen, Cell reports 34 (2021).

106 G. Tesei and K. Lindorff-Larsen, Open Research Europe 2, 94 (2023).

107 J. A. Joseph, A. Reinhardt, A. Aguirre, P. Y. Chew, K. O. Russell, J. R. Espinosa, A. Garaizar, and R. Collepardo-Guevara, Nature Computational Science 1, 732 (2021).

108 A. R. Tejedor, A. Aguirre Gonzalez, M. J. Maristany, P. Y. Chew, K. Russell, J. Ramirez, J. R. Espinosa, and R. Collepardo-Guevara, ACS Central Science 11, 302 (2025).

109 S. Liu, C. Wang, and B. Zhang, Biochemistry (2025).

110 S. Liu, C. Wang, A. P. Latham, X. Ding, and B. Zhang, PLoS Computational Biology 19, e1011442 (2023).

111 A. R. Tejedor, J. R. Tejedor, and J. Ramírez, The Journal of Chemical Physics 157 (2022).

112 A. R. Tejedor, R. Collepardo-Guevara, J. Ramirez, and J. R. Espinosa, The Journal of Physical Chemistry B 127, 4441 (2023).

113 D. Sundaravadivelu Devarajan, J. Wang, B. Szala-Mendyk, S. Rekhi, A. Nikoubashman, Y. C. Kim, and J. Mittal, Nature Communications 15, 1912 (2024).

114 D. Sundaravadivelu Devarajan and J. Mittal, JACS Au 4, 4394 (2024).

115 A. Ladd and L. Woodcock, Chemical Physics Letters 51, 155 (1977).

116 J. R. Espinosa, J. A. Joseph, I. Sanchez-Burgos, A. Garaizar, D. Frenkel, and R. Collepardo-Guevara, Proceedings of the National Academy of Sciences 117, 13238 (2020).

117 C. Vega, E. Sanz, J. L. F. Abascal, and E. G. Noya, Journal of Physics: Condensed Matter 20, 153101 (2008).

118 D. A. Kofke, The Journal of Chemical Physics 98, 4149 (1993).

119 D. A. Kofke, Molecular Physics 78, 1331 (1993).

120 S. P. Somasekharan, A. El-Naggar, G. Leprivier, H. Cheng, S. Hajee, T. G. Grunewald, F. Zhang, T. Ng, O. Delattre, V. Evdokimova, et al., Journal of Cell Biology 208, 913 (2015).

121 B. S. Schuster, G. L. Dignon, W. S. Tang, F. M. Kelley, A. K. Ranganath, C. N. Jahnke, A. G. Simpkins, R. M. Regy, D. A. Hammer, M. C. Good, et al., Proceedings of the National Academy of Sciences 117, 11421 (2020).

122 X. Wang, S. Ramírez-Hinestrosa, J. Dobnikar, and D. Frenkel, Physical Chemistry Chemical Physics 22, 10624 (2020).

123 E. Pedraza, D. Hoyos, A. Feito, F. Gámez, I. Sanchez-Burgos, R. Collepardo-Guevara, A. R. Tejedor, and J. R. Espinosa, bioRxiv (2025), 10.1101/2025.03.26.645197.

124 J. S. Rowlinson and B. Widom, Molecular theory of capillarity (Courier Corporation, 2013).

125 C. Pak, M. Kosno, A. Holehouse, S. Padrick, A. Mittal, R. Ali, A. Yunus, D. Liu, R. Pappu, and M. Rosen, Molecular Cell 63, 72 (2016).

126 W. Borcherds, A. Bremer, M. B. Borgia, and T. Mittag, Current Opinion in Structural Biology 67, 41 (2021).

127 J. Wang, J.-M. Choi, A. S. Holehouse, H. O. Lee, X. Zhang, M. Jahnel, S. Maharana, R. Lemaitre, A. Pozniakovsky, D. Drechsel, I. Poser, R. V. Pappu, S. Alberti, and A. A. Hyman, Cell 174, 688 (2018).

128 I. Sanchez-Burgos, A. R. Tejedor, R. Collepardo-Guevara, J. B. de la Serna, and J. R. Espinosa, bioRxiv, 2025.03.11.642656 (2025).

129 P. R. Banerjee, A. N. Milin, M. M. Moosa, P. L. Onuchic, and A. A. Deniz, Angewandte Chemie 129, 11512 (2017).

130 N. H Espejo, A. Feito, I. Sanchez-Burgos, A. Garaizar, M. M. Conde, A. Rey, A. Castro, R. Collepardo-Guevara, A. R. Tejedor, and J. R Espinosa, bioRxiv, 2025.02.21.639421 (2025).

131 R. S. Fisher and S. Elbaum-Garfinkle, Nature Communications 11, 1 (2020).

132 P. Yang, C. Mathieu, R.-M. Kolaitis, P. Zhang, J. Messing, U. Yurtsever, Z. Yang, J. Wu, Y. Li, Q. Pan, J. Yu, E. W. Martin, T. Mittag, H. J. Kim, and J. P. Taylor, Cell 181, 325 (2020).

133 X.-M. Liu, L. Ma, R. Schekman, S. R. Pfeffer, and Y. G. Zhao, eLife 10, e71982 (2021).

134 K. A. Burke, A. M. Janke, C. L. Rhine, and N. L. Fawzi, Molecular Cell 60, 231 (2015).

135 J. P. Brady, P. J. Farber, A. Sekhar, Y.-H. Lin, R. Huang, A. Bah, T. J. Nott, H. S. Chan, A. J. Baldwin, J. D. Forman-Kay, and L. E. Kay, Proceedings of the National Academy of Sciences 114, E8194 (2017).

136 S. Ambadipudi, J. Biernat, D. Riedel, E. Mandelkow, and M. Zweckstetter, Nature Communications 8, 1 (2017).

137 H. Deng, K. Gao, and J. Jankovic, Nature Reviews Neurology 10, 337 (2014).

138 P. Yang, C. Mathieu, R.-M. Kolaitis, P. Zhang, J. Messing, U. Yurtsever, Z. Yang, J. Wu, Y. Li, Q. Pan, et al., Cell 181, 325 (2020).

139 A. G. Niaki, J. Sarkar, X. Cai, K. Rhine, V. Vidaurre, B. Guy, M. Hurst, J. C. Lee, H. R. Koh, L. Guo, et al., Molecular Cell 77, 82 (2020).

140 M. Linsenmeier, L. Faltova, C. Morelli, U. Capasso Palmiero, C. Seiffert, A. M. Küffner, D. Pinotsi, J. Zhou, R. Mezzenga, and P. Arosio, Nature Chemistry 15, 1340 (2023).

141 N. A. Erkamp, I. Sanchez-Burgos, A. Zhou, T. J. Krug, S. Qamar, T. Sneideris, E. Zhang, K. Nakajima, A. Chen, R. Collepardo-Guevara, J. van Hest, P. St George-Hyslop, D. A. Weitz, J. R. Espinosa, and T. P. J. Knowles, bioRxiv, 2024.11.15.623768 (2024).

142 J. Wessén, S. Das, T. Pal, and H. S. Chan, The Journal of Physical Chemistry B 126, 9222 (2022).

143 G. L. Dignon, W. Zheng, and J. Mittal, Current opinion in chemical engineering 23, 92 (2019).

144 R. M. Regy, J. Thompson, Y. C. Kim, and J. Mittal, Protein Science 30, 1371 (2021).

145 T. Dannenhoffer-Lafage and R. B. Best, The Journal of Physical Chemistry B 125, 4046 (2021).

146 J. M. Lotthammer, G. M. Ginell, D. Griffith, R. Emenecker, and A. S. Holehouse, Biophysical Journal 123, 43a (2024).

147 I. Sanchez-Burgos, L. Herriott, R. Collepardo-Guevara, and J. R. Espinosa, Biophysical Journal 122, 2973 (2023).

148 R. Haider, S. Penumutchu, S. Boyko, and W. K. Surewicz, Biophysical Journal 123, 361 (2024).

149 D. Frenkel and B. Smit, Understanding Molecular Simulation: From Algorithms to Applications, 3rd ed. (Academic Press, San Diego, CA, 2023).

