## Supplementary Material for "Predicting Saturation Concentrations of Phase-Separating Proteins via Thermodynamic Integration"

Eduardo Pedraza†

*Department of Physical Chemistry, Universidad Complutense de Madrid, Av. Complutense s/n, Madrid 28040, Spain †These authors contributed equally*

Andrés R. Tejedor†

*Yusuf Hamied Department of Chemistry, University of Cambridge, Lensfield Road, Cambridge CB2 1EW, UK and Department of Physical Chemistry, Universidad Complutense de Madrid, Av. Complutense s/n, Madrid 28040, Spain †These authors contributed equally*

Alejandro Feito, Francisco Gámez, and Eduardo Sanz*

*Department of Physical Chemistry, Universidad Complutense de Madrid, Av. Complutense s/n, Madrid 28040, Spain*

Rosana Collepardo-Guevara

*Yusuf Hamied Department of Chemistry, University of Cambridge, Lensfield Road, Cambridge CB2 1EW, UK and Department of Genetics, University of Cambridge, Cambridge, UK*

Jorge R. Espinosa†

*Department of Physical Chemistry, Universidad Complutense de Madrid, Av. Complutense s/n, Madrid 28040, Spain and Yusuf Hamied Department of Chemistry, University of Cambridge, Lensfield Road, Cambridge CB2 1EW, UK*

(Dated: May 9, 2025)

### SI. THE MPIPI-RECHARGED MODEL

We use the Mpipi-Recharged model, a residue-level coarse-grained force field for both protein and RNA condensates [1]. In this model, each amino acid or nucleotide is represented by a single bead connected by harmonic bonds. Globular domains are treated as rigid bodies whose beads are fixed at the $C_\alpha$ of the corresponding Protein Data Bank (PDB). The potential energy is computed as the sum of pairwise bonded ($E_{\text{bonded}}$) and non-bonded ($E_{\text{non-bonded}}$) interactions as:

$$E = E_{\text{bonded}} + E_{\text{non-bonded}}. \tag{S1}$$

The intrinsically disordered regions (IDRs) are modelled as fully flexible polymers and these are connected to the globular domains. The bonded potential is written as:

$$E_{\text{bonded}}(r_{ij}) = \sum_{ij} k(r_{ij} - r_0)^2, \tag{S2}$$

where $r_{ij}$ is the distance between the connected beads, $r_0 = 3.81\,\text{Å}$ is the equilibrium bond length, and the spring constant $k = 9.6\,\text{kcal} \cdot \text{mol}^{-1} \cdot \text{Å}^{-2}$. The sum runs over all paired amino acids.

Non-bonded interactions consist of the sum of the hydrophobic interaction and electrostatic interaction. The hydrophobic interaction is given by the Wang–Frenkel (WF) potential [2] that accounts for short-ranged excluded-volume repulsion and long-ranged attraction. This potential is defined as

$$E_{\text{WF}}(r_{i,j}) = \sum_{ij} \epsilon_{ij}\alpha_{ij} \left[\left(\frac{\sigma_{ij}}{r_{ij}}\right)^{2\mu_{ij}} - 1\right]\left[\left(\frac{R_{ij}}{r_{ij}}\right)^{2\mu_{ij}} - 1\right]^{2\nu_{ij}}, \tag{S3}$$

where

$$\alpha_{ij} = 2\nu_{ij}\left(\frac{R_{ij}}{\sigma_{ij}}\right)^{2\mu_{ij}} \left\{\frac{2\nu_{ij}+1}{2\nu_{ij}\left[\left(\frac{R_{ij}}{\sigma_{ij}}\right)^{2\mu_{ij}} - 1\right]}\right\}^{2\nu_{ij}+1}. \tag{S4}$$

Here $\sigma_{ij}$ is the pair-of-beads diameter, defined from the individual diameter ($\sigma_i$ and $\sigma_j$) assuming the Lorentz-Berthelot mixing rules (i.e., $\sigma_{ij} = (\sigma_i + \sigma_j)/2$). $R_{ij} = 3\sigma_{ij}$ is the cut-off distance for the $ij$-th interaction. The interaction parameter $\epsilon_{ij}$ is defined for each specific amino acid pair based on our atomistic Potential of Mean Force calculations and

*
†

bioinformatics data [1]. The exponent $\mu_{ij}$ is set to 1 for all the pairs and $\nu_{ij}$ depends on the specific pair, ranging from 2 to 12 (see Ref. [1]). Notice that higher values of $\mu_{ij}$ lead to a steeper increase in the repulsive part of the potential.

The electrostatic interactions are described by the Yukawa potential [3] instead of the Debye–Hückel potential [4] originally employed in the Mpipi model. Avoiding the use of explicit charge values, this potential allows to modulate independently the strength of electrostatic interactions in a pair-specific basis. This potential is defined as

$$E_{\text{electrostatic}} = \sum_{ij} \frac{A_{ij}}{r_{ij}} \exp(-\kappa r_{ij}), \qquad r_{ij} < r_c, \tag{S5}$$

where $A_{ij}$ is the interaction parameter, $\kappa$ is the screening due to ions, and $r_{ij}$ is the distance between interacting pairs. The cut-off for the electrostatic interaction, $r_c$, is set to 3.5 nm. The screening parameter $\kappa$ is expressed in a explicit way as $\kappa = \sqrt{8\pi B c_s}$, where $c_s$ is the salt concentration—normally set to 150 mM of NaCl—and $B = e_0^2/4\pi k_B T \varepsilon_0 \varepsilon_r$ is the Bjerrum length. The relative dielectric constant $\varepsilon_r$ varies with temperature according to the empiric formula [5]:

$$\varepsilon_r(T) = \frac{5321}{T} + 233.760 - 0.9297T + 1.417 \cdot 10^{-3} T^2 - 8.292 \cdot 10^{-7} T^3, \tag{S6}$$

for $T$ in Kelvin. The pair-specific optimized values of $A_{ij}$ of the Yukawa potential can be found in Ref. [1]. In the Mpipi-Recharged model, such parameters indicate that the interaction between oppositely charged pairs is significantly stronger than those between identically charged pairs.

### SII. CALCULATION OF THE PHASE DIAGRAM VIA DIRECT COEXISTENCE SIMULATIONS

The determination of the phase diagrams in the temperature-density plane was performed using the Direct Coexistence method [6, 7]. The proteins are placed in a prismatic elongated box to simulate both the high-density and low-density phases separated by an interface. The long side of the box is perpendicular to the interfaces. Since determination of the optimal box dimensions is key to minimising finite size effects while ensuring computer efficiency, we provide the following guidelines

1. A minimum of 48 protein replicas for proteins with globular regions and a minimum of 100 protein replicas for flexible protein systems per box were used.

2. To avoid self-interactions through periodic boundary conditions we enforce the short sides of the box to be larger than at least twice the radius of gyration of the protein in the system.

3. The long side of the box should keep the total density of the system at approximately $\sim 0.1\,\mathrm{g{\cdot}cm^{-3}}$.

Simulations are carried out in the canonical (NVT) ensemble using a Nosé-Hover thermostat [8] for the rigid bodies (representing the globular domains) integrated in the RIGID package, and a Langevin thermostat [9] for the particles in flexible regions, both with a relaxation time of 5 ps. The timestep for the Verlet integration of the equations of motion is 10 fs. After an equilibration period ($\sim$ 10ns), we run the simulation of 1-2 $\mu$s depending on the specific system. If two different phases are detectable—i.e., a high density phase and a low density phase—, the densities are calculated.

The critical density ($\rho_c$) and temperature ($T_c$) of the phase diagrams are evaluated by means of the law of rectilinear diameters and critical exponents [10, 11]:

$$\frac{\rho_l(T) + \rho_d(T)}{2} = \rho_c + s_2(T_c - T), \tag{S7}$$

and

$$(\rho_l(T) - \rho_d(T))^{\beta} = d\left(1 - \frac{T}{T_c}\right), \tag{S8}$$

where the critical exponent $\beta = 3.06$, $\rho_d$ and $\rho_l$ are the coexisting densities of the diluted and condensed phases, respectively, and $d$ and $s_2$ are fitting parameters.

### SIII. SIMULATION DETAILS TO OBTAIN THE INTERNAL ENERGIES

The internal energies used for the TI-$C_{sat}$ and TI-$CC$ Methods were estimated as the sum of the kinetic energy and the pairwise interaction energy. All energy values were obtained from simulations performed in the NVT ensemble. A minimum of 48 protein replicas for proteins with globular regions and a minimum of 100 protein replicas for flexible protein systems per box were used. All NVT simulations were carried out using the molecular dynamics LAMMPS software [12], version 2nd of August of 2023. All simulations use a Nosé-Hover thermostat [8] for the rigid bodies (representing the globular domains) integrated in the RIGID package, and a Langevin thermostat [9] for the particles in flexible regions, both with a relaxation time of 5 ps. The timestep for the Verlet integration of the equations of motion is 10 fs. After equilibration has reached, we run the simulation of 1 $\mu$s. The cut-off values used for the interactions are those described in Section SI.

### SIV. SEQUENCES USED IN THIS WORK

#### FUS

MASNDYTQQATQSYGAYPTQPGQGYSQQSSQPYGQQSYSGYSQSTDTSGYGQSSYSSYGQSQNTG
YGTQSTPQGYGSTGGYGSSQSSQSSYGQQSSYPGYGQQPAPSSTSGSYGSSSQSSSYGQPQSGSYSQ
QPSYGGQQQSYGQQQSYNPPQGYGQQNQYNSSSGGGGGGGGGGNYGQDQSSMSSGGGSGGGYG
NQDQSGGGGSGGYGQQDRGGRGRGGSGGGGGGGGGGYNRSSGGYEPRGRGGGRGGRGGMGGS
DRGGFNKFGGPRDQGSRHDSEQDNSDNNTIFVQGLGENVTIESVADYFKQIGIIKTNKKTGQPMIN
LYTDRETGKLKGEATVSFDDPPSAKAAIDWFDGKEFSGNPIKVSFATRRADFNRGGGNGRGGRGR
GGPMGRGGYGGGGSGGGGRGGFPSGGGGGGGQQRAGDWKCPNPTCENMNFSWRNECNQCKA
PKPDGPGGGPGGSHMGGNYGDDRRGGRGGYDRGGYRGRGGDRGGFRGGRGGGDRGGFGPGK
MDSRGEHRQDRRERPY

We have used two PDB codes to simulate the globular regions of FUS: residues from 285–371 (PDB code: 2LCW) and from 422–453 (PDB code: 6G99).

#### FUS-LCD

MASNDYTQQATQSYGAYPTQPGQGYSQQSSQPYGQQSYSGYSQSTDTSGYGQSSYSSYGQSQNTG
YGTQSTPQGYGSTGGYGSSQSSQSSYGQQSSYPGYGQQPAPSSTSGSYGSSSQSSSYGQPQSGSYSQ
QPSYGGQQQSYGQQQSYNPPQGYGQQNQYNS

#### hnRNPA1

MSKSESPKEPEQLRKLFIGGLSFETTDESLRSHFEQWGTLTDCVVMRDPNTKRSRGFGFVTYATVE
EVDAAMNARPHKVDGRVVEPKRAVSREDSQRPGAHLTVKKIFVGGIKEDTEEHHLRDYFEQYGKI
EVIEIMTDRGSGKKRGFAFVTFDDHDSVDKIVIQKYHTVNGHNCEVRKALSKQEMASASSSQRGRS

GSGNFGGGRGGGFGGNDNFGRGGNFSGRGGFGGSRGGGGYGGSGDGYNGFGNDGGYGGGGPG
YSGGSRGYGSGGQGYGNQGSGYGGSGSYDSYNNGGGGGFGGGSGSNFGGGGSYNDFGNYNNQSS
NFGPMKGGNFGGRSSGPYGGGGQYFAKPRNQGGYGGSSSSSSYGSGRRF

We have used the PDB code to simulate two globular regions of hnRNPA1, residues from 9–91 and from 103–181, both in the same PDB (PDB code: 1L3K).

**hnRNPA1-LCD**

GSMASASSSQRGRSGSGNFGGGRGGGFGGNDNFGRGGNFSGRGGFGGSRGGGGYGGSGDGYNGF
GNDGSNFGGGGSYNDFGNYNNQSSNFGPMKGGNFGGRSSGPYGGGGQYFAKPRNQGGYGGSSSS
SSYGSGRRF

**TDP43**

MSEYIRVTEDENDEPIEIPSEDDGTVLLSTVTAQFPGACGLRYRNPVSQCMRGVRLVEGILHAPDAG
WGNLVYVVNYPKDNKRKMDETDASSAVKVKRAVQKTSDLIVLGLPWKTTEQDLKEYFSTFGEVL
MVQVKKDLKTGHSKGFGFVRFTEYETQVKVMSQRHMIDGRWCDCKLPNSKQSQDEPLRSRKVFV
GRCTEDMTEDELREFFSQYGDVMDVFIPKPFRAFAFVTFADDQIAQSLCGEDLIIKGISVHISNAEPK
HNSNRQLERSGRFGGNPGGFGNQGGFGNSRGGGAGLGNNQGSNMGGGMNFGAFSINPAMMAAA
QAALQSSWGMMGMLASQQNQSGPSGNNQNQGNMQREPNQAFGSGNNSYSGSNSGAAIGWGSASN
AGSGSGFNGGFGSSMDSKSSGWGM

We have used several PDB codes to simulate the globular regions of TDP43: residues from 2-38, 40-49, 51-79 (PDB code: 5MDI), residues from 103-179 (PDB code: 2CQG), residues from 193-267 (PDB code: 1WF0) and for the residues 307-349 (PDB code: 2N2C). Moreover, for the globular region of residues 307–349, we increased the $\epsilon_{ij}$ values corresponding to interactions within this same region by 10%.

**G3BP1**

MVMEKPSPLLVGREFVRQYYTLLNQAPDMLHRFYGKNSSYVHGGLDSNGKPADAVYGQKEIHRK
VMSQNFTNCHTKIRHVDAHATLNDGVVVQVMGLLSNNNQALRRFMQTFVLAPEGSVANKFYVHN
DIFRYQDEVFGGFVTEPQEESEEEVEEPEERQQTPEVVPDDSGTFYDQAVVSNDMEEHLEEPVAE
PEPDPEPEPEQEPVSEIQEEKPEPVLEETAPEDAQKSSSPAPADIAQTVQEDLRTFSWASVTSKNLP

PSGAVPVTGIPPHVVKVPASQPRPESKPESQIPPQRPQRDQRVREQRINIPPQRGPRPIREAGEQGD
IEPRRMVRHPDSHQLFIGNLPHEVDKSELKDFFQSYGNVVELRINSGGKLPNFGFVVFDDSEPVQKV
LSNRPIMFRGEVRLNVEEKKTRAAREGDRRDNRLRGPGGPRGGLGGGMRGPPRGGMVQKPGFG
VGRGLAPRQMVMEKPSPLLVGREFVRQYYTLLNQAPDMLHRFYGKNSSYVHGGLDSNGKPADAV
YGQKEIHRKVMSQNFTNCHTKIRHVDAHATLNDGVVVQVMGLLSNNNQALRRFMQTFVLAPEGS
VANKFYVHNDIFRYQDEVFGGFVTEPQEESEEEVEEPEERQQTPEVVPDDSGTFYDQAVVSNDME
EHLEEPVAEPEPDPEPEPEQEPVSEIQEEKPEPVLEETAPEDAQKSSSPAPADIAQTVQEDLRTFSW
ASVTSKNLPPSGAVPVTGIPPHVVKVPASQPRPESKPESQIPPQRPQRDQRVREQRINIPPQRGPRP
IREAGEQGDIEPRRMVRHPDSHQLFIGNLPHEVDKSELKDFFQSYGNVVELRINSGGKLPNFGFVVF
DDSEPVQKVLSNRPIMFRGEVRLNVEEKKTRAAREGDRRDNRLRGPGGPRGGLGGGMRGPPRGG
MVQKPGFGVGRGLAPRQ

We have used one PDB code to simulate the globular region of the dimerization domain of G3BP1: residues from 7-41, 52-117, 124-137, 473-507, 518-583, 590-603 (PDB code:3Q90), and we have used AlphaFold for their RRM domains: residues from 338-413 and 804-879 with a confidence level of 90%.

**YBX1**

MSSEAETQQPPAAPPAAPALSAADTKPGTTGSGAGSGGPGGLTSAAPAGGDKKVIATKVLGTVK
WFNVRNGYGFINRNDTKEDVFVHQTAIKKNNPRKYLRSVGDGETVEFDVVEGEKGAEAANVTGP
GGVPVQGSKYAADRNHYRRYPRRRGPPRNYQQNYQNSESGEKNEGSESAPEGQAQQRRPYRRRR
FPPYYMRRPYGRRPQYSNPPVQGEVMEGADNQGAGEQGRPVRQNMYRGYRPRFRRGPPRQRQP
REDGNEEDKENQGDETQGQQPPQRRYRRNFNYRRRRPENPKPQDGKETKAADPPAENSSAPEAE
QGGAE

We have used AlphaFold to simulate the globular regions of YBX1: residues from 52–67, from 72-77, from 83-87, from 108-116 and from 119-133 with a confidence level of 90%.

**LAF-1-RGG**

MESNQSNNGGSGNAALNRGGRYVPPHLRGGDGGAAAAASAGGDDRRGGAGGGGYRRGGGNSGG
GGGGGYDRGYNDNRDDRDNRGGSGGYGRDRNYEDRGYNGGGGGGGNRGYNNNRGGGGGGYN
RQDRGDGGSSNFSRGGYNNRDEGSDNRGSGRSYNNDRRDNGGDGLEHHHHHH

#### DDX4-wt

MGDEDWEAEINPHMSSYVPIFEKDRYSGENGDNFNRTPASSSEMDDGPSRRDHFMKSGFASGRNF
GNRDAGECNKRDNTSTMGGFGVGKSFGNRGFSNSRFEDGDSSGFWRESSNDCEDNPTRNRGFSK
RGGYRDGN

#### DDX4-cs

MGDRDWRAEINPHMSSYVPIFEKDRYSGENGRNFNDTPASSSEMRDGPSERDHFMKSGFASGDNF
GNRDAGKCNERDNTSTMGGFGVGKSFGNEGFSNSRFERGDSSGFWRESSNDCRDNPTRNDGFSDR
GGYEKGN

#### hnRNPA1-LCD variants

#### -12F+12Y (allY)

GSMASASSSQRGRSGSGNYGGGRGGGYGGNDNYGRGGNYSGRGGYGGSRGGGGYGGSGDGYNG
YGNDGSNYGGGGSYNDYGNYNNQSSNYGPMKGGNYGGRSSGGSGGGGQYYAKPRNQGGYGGSS
SSSSYGSGRRY

#### +7Y-7F (allF)

GSMASASSSQRGRSGSGNFGGGRGGGFGGNDNFGRGGNFSGRGGFGGSRGGGGFGGSGDGFNGF
GNDGSNFGGGGSFNDFGNFNNQSSNFGPMKGGNFGGRSSGGSGGGGQFFAKPRNQGGFGGSSSSS
SFGSGRRF

#### -12F+12W (allW)

GSMASASSSQRGRSGSGNWGGGRGGGWGGNDNWGRGGNWSGRGGWGGSRGGGGWGGSGDG
WNGWGNDGSNWGGGGSWNDWGNWNNQSSNWGPMKGGNWGGRSSGSGGGGGQWWAKPRNQ
GGWGGSSSSSSWGSGRRW

### -6R

GSMASASSSQGGRSGSGNFGGGRGGGFGGNDNFGGGGNFSGSGGFGGSRGGGGYGGSGDGYNGF
GNDGSNFGGGGSYNDFGNYNNQSSNFGPMKGGNFGGSSSGPYGGGGQYFAKPGNQGGYGGSSSS

SSYGSGGRF

**-3R+3K**

GSMASASSSQRGKSGSGNFGGGRGGGFGGNDNFGRGGNFSGRGGFGGSKGGGGYGGSGDGYNGF
GNDGSNFGGGGSYNDFGNYNNQSSNFGPMKGGNFGGRSSGGSGGGGQYFAKPRNQGGYGGSSSS
SSYGSGRKF

**+8D**

GSMASASSSQRDRSGSGNFGGGRDGGFGGNDNFGRGDNFSGRGDFGGSRDGGGYGGSGDGYNGF
GNDGSNFGGGGSYNDFGNYNNQSSNFGPMKGGNFGGRSSDPYGGGGQYFAKPRNQDGYGGSSSS
SSYDSGRRF

**+12E**

GSMASAESSQREREESGNFGEGRGGGFGGNDNFGRGGNFSERGGFGGSRGEGGYGGEGDGYNGF
GNDGSNFGGGGSYNDFGNYNNQSSNFEPMKGGNFGERSSGPYEGGGQYFAKPRNQGGYGGSSSSS
SYGSERRF

**+12D**

GSMASADSSQRDRDDSGNFGDGRGGGFGGNDNFGRGGNFSDRGGFGGSRGDGGYGGDGDGYNG
FGNDGSNFGGGGSYNDFGNYNNQSSNFDPMKGGNFGDRSSGPYDGGGQYFAKPRNQGGYGGSSS
SSSYGSDRRF

**-4D**

GSMASASSSQRGRSGSGNFGGGRGGGFGGNGNFGRGGNFSGRGGFGGSRGGGGYGGSGGGYNGF
GNSGSNFGGGGSYNGFGNYNNQSSNFGPMKGGNFGGRSSGPYGGGGQYFAKPRNQGGYGGSSSSS
SYGSGRRF

**+7R**

GSMASASSSQRGRSGRGNFGGGRGGGFGGNDNFGRGGNFSGRGGFGGSRGGGRYGGSGDRYNGF
GNDGRNFGGGGSYNDFGNYNNQSSNFGPMKGGNFRGRSSGPYGRGGQYFAKPRNQGGYGGSSSS
RSYGSGRRF

**-9F+6Y**

GSMASASSSQRGRSGSGNFGGGRGGGYGGNDNYGRGGNYSGRGGFGGSRGGGGYGGSGDGYNG
GGNDGSNYGGGGSYNDSGNYNNQSSNFGPMKGGNYGGRSSGGSGGGGQYGAKPRNQGGYGGSSS
SSSYGSGRRY

**-8F+4Y**

GSMASASSSQRGRSGSGNFGGGRGGGYGGNDNGGRGGNYSGRGGFGGSRGGGGYGGSGDGYNG
GGNDGSNYGGGGSYNDSGNYNNQSSNFGPMKGGNYGGRSSGGSGGGGQYGAKPRNQGGYGGSSS
SSSYGSGRRF

**+7R+12D**

GSMASADSSQRDRDDRGNFGDGRGGGFGGNDNFGRGGNFSDRGGFGGSRGDGRYGGDGDRYNG
FGNDGRNFGGGGSYNDFGNYNNQSSNFDPMKGGNFRDRSSGPYDRGGQYFAKPRNQGGYGGSSS
SRSYGSDRRF

**+7K+12D**

GSMASADSSQRDRDDKGNFGDGRGGGFGGNDNFGRGGNFSDRGGFGGSRGDGKYGGDGDKYNG
FGNDGKNFGGGGSYNDFGNYNNQSSNFDPMKGGNFKDRSSGPYDKGGQYFAKPRNQGGYGGSSS
SKSYGSDRRF

### SV. SENSITIVITY OF THE RESULTS TO THE INTEGRATION DENSITY

To evaluate the sensitivity of the results to the coexistence densities at which the thermodynamic integration is performed, the allF sequence was used. First, the saturation concentrations were determined at the coexistence densities indicated by the phase diagram at the temperature closest to the critical point. Then, the saturation concentrations were calculated again using the same coexistence density for the dense phase as in the previous case, but assigning an arbitrary coexistence density for the dilute phase. The results, shown in Fig. S1, reveal a marked dependence of the obtained values on the coexistence density used in the thermodynamic integration.

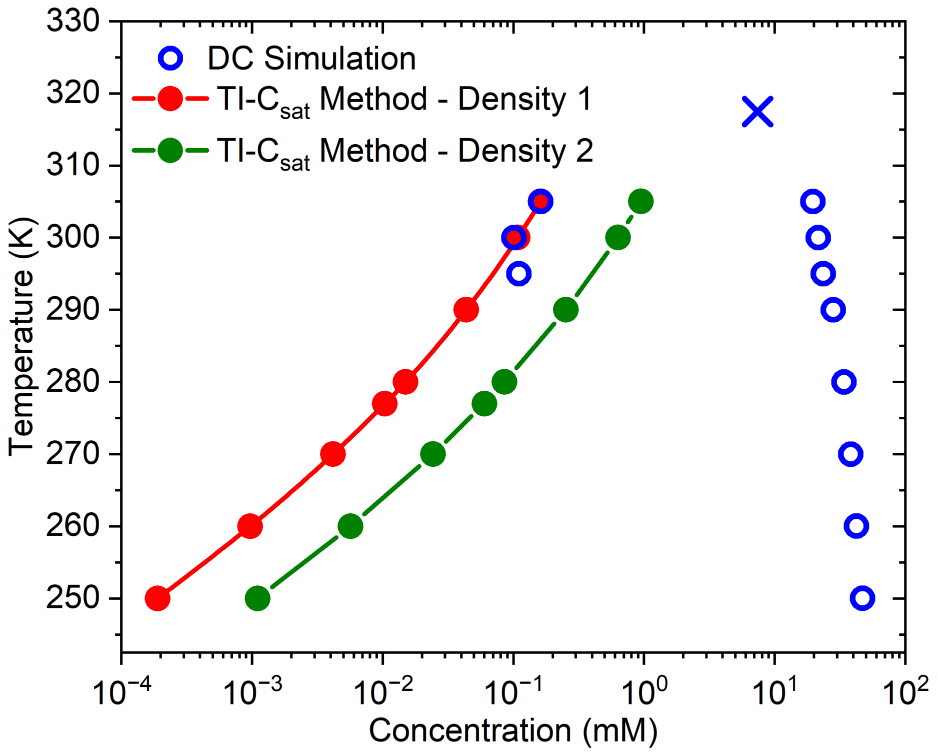


**FIG. S1:** Sensitivity of the results to the density at which the integration is performed. Phase diagrams (empty symbols) of allF in the temperature-density plane obtained from DC simulations using the Mpipi-Recharged models. Red and green symbols represent the saturation concentrations ($C_{sat}$) at different temperatures calculated via TI-$C_{sat}$ starting from the closest temperature to the critical point at two different dilute densities.

### SVI. COMPARISON OF NORMALIZED PHASE DIAGRAMS

In Fig. S2, we present the temperature–concentration phase diagrams for the studied proteins, renormalised by their own critical temperature and critical density. It can be observed that the resulting phase diagrams clearly differ in the dilute branch for globular proteins (Fig. S2.a), highlighting the non-trivial behaviour of low-density phase. In the case of the hnRNPA1 variants (Fig. S2.b), the differences are more subtle. This is reasonable considering that the saturation concentration at the critical point is similar across all variants, and consistently falling within the range of $(6.73, 10.31)$ mM.

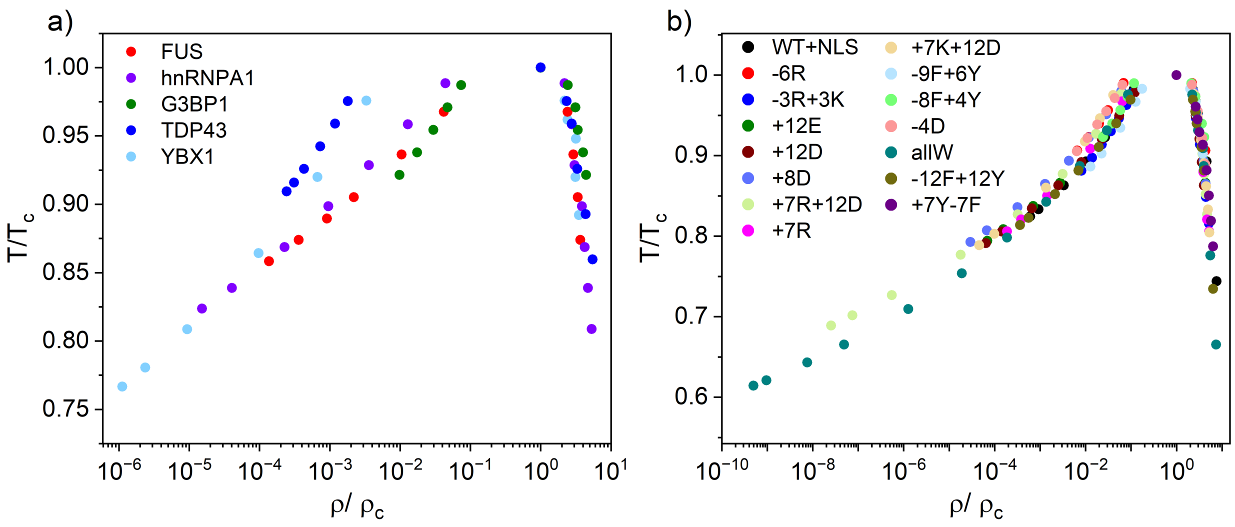


**FIG. S2:** Temperature-concentration phase diagram for globular proteins (a) and flexible proteins (b) where both temperature and density are renormalized by their own critical temperature and critical density.

### SVII. CHARGE DISTRIBUTION IN HNRNPA1 VARIANTS SEQUENCES

The Mpipi-Recharged model employs an implicit solvent and electrolyte, which constitutes an approximation of the physical conditions of the system and it can mitigate some of the limitations associated with the use of an implicit solvent and electrolyte [1]. We investigated whether the distribution of charged residues along the sequences is responsible for the observed deviations. To this end, we used mutant sequences of the low-complexity domain of hnRNPA1. Figure S3 displays the diagrams of the sequences, indicating the charge character of each residue and its position within the hnRNPA1-LCD variants used. It can be observed that in some variants, such as +7R+12D, +7K+12D, or +7R, where the model overestimates the critical temperature ($T_c$) charged residues are abundant.

Furthermore, we calculate the blockiness parameter ($B_{LC}$) [13] to assess the influence of the charge distribution of the sequences. Charge blockiness is calculated as the sum of the charges (R and K, +1; E and D, −1) using a sliding window of 25 residues centred on each residue of the sequence. For residues at the termini, the sum was calculated using the available amino acids. For example, for residue 3 in the sequence, the charge was determined as the sum of the charges from residues 1 to 15. In addition, the blockiness parameter was calculated using the following formula [13]:

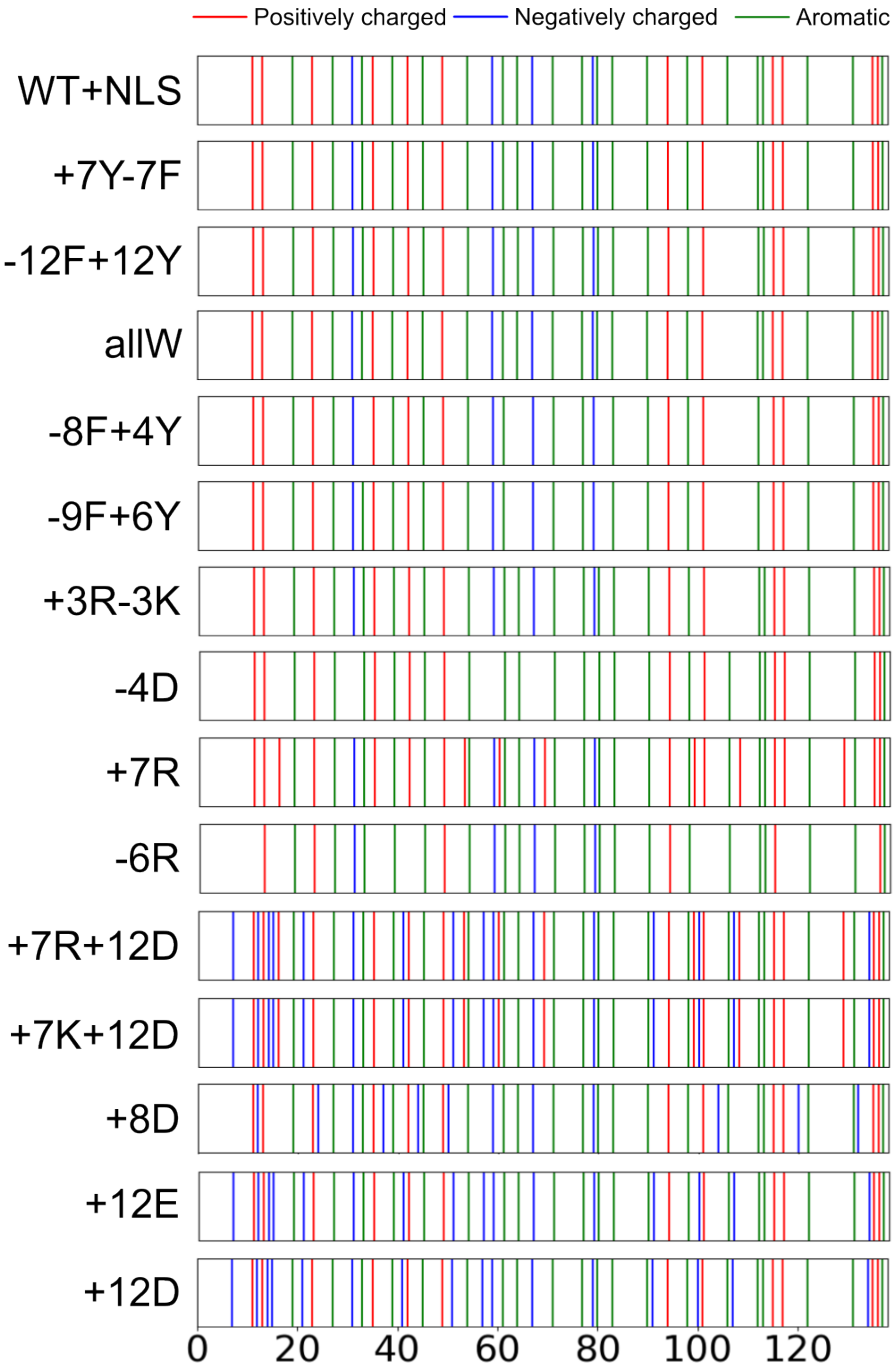


**FIG. S3:** Diagram of content type and placement in hnRNPA1-LCD variants.

$$B_{\rm LC} = \left(C_{\max(+)} + C_{\max(-)}\right) \cdot \frac{\sum_{k=1}^{Nd(+,-)} d_k(+,-)}{Nd(+,-)} \Big/ \left( \frac{\sum_{k=1}^{Nd(+,+)} d_k(+,+)}{Nd(+,+)} + \frac{\sum_{k=1}^{Nd(-,-)} d_k(-,-)}{Nd(-,-)} \right) \tag{S9}$$

where $C_{\max(+)}$ and $C_{\max(-)}$ are the absolute maximum positive and negative values in the charge plot, respectively, $d(+,+)$, $d(-,-)$ and $d(+,-)$ are the distances between pairs of charged residues positive (+), or negative (−), and $Nd(+,+)$, $Nd(-,-)$, $Nd(+,-)$ represents the number of these pairs.

In Fig. S4, the charge plots of the variants WT+NLS, -12F+12Y, +7R+12D, +7R, +8D and +12D are shown, together with their corresponding calculated $B_{LC}$ values. However, no significant differences are observed that would account for the model overestimations based on charge blockiness within the sequences.

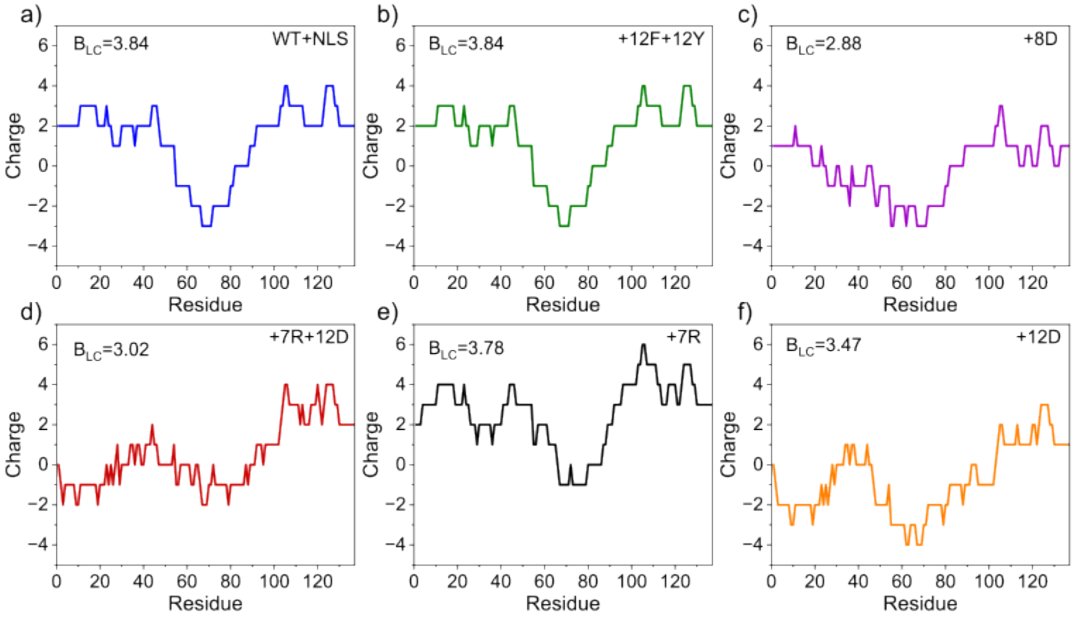


**FIG. S4:** Charge plots for different hnRNPA1 mutants with their $B_{LC}$ parameter.

### SVIII. DEVIATION FROM THE IDEAL LINE IN INTERPOLATIONS

The values of the metric, $D$, quantify the average deviation between the computational predictions and the experimental values. The following formula is used to calculate the parameter $D$

$$D = \frac{1}{n}\sum_{i=1}^{n}(x_{exp,i} - x_{sim,i})^2 \tag{S10}$$

[1] A. R. Tejedor, A. Aguirre Gonzalez, M. J. Maristany, P. Y. Chew, K. Russell, J. Ramirez, J. R. Espinosa, and R. Collepardo-Guevara, “Chemically Informed Coarse-Graining of Electrostatic Forces in Charge-Rich Biomolecular Condensates,” *ACS Central Science*, vol. 11, pp. 302–321, Feb. 2025.

[2] X. Wang, S. Ramírez-Hinestrosa, J. Dobnikar, and D. Frenkel, “The lennard-jones potential: when (not) to use it,” *Physical Chemistry Chemical Physics*, vol. 22, no. 19, pp. 10624–10633, 2020.

[3] H. Yukawa, “On the interaction of elementary particles. i,” *Proceedings of the Physico-Mathematical Society of Japan. 3rd Series*, vol. 17, pp. 48–57, 1935.

[4] P. Debye and E. Hückel, “De la theorie des electrolytes. i. abaissement du point de congelation et phenomenes associes,” *Physikalische Zeitschrift*, vol. 24, no. 9, pp. 185–206, 1923.

[5] G. Akerlof and H. Oshry, “The dielectric constant of water at high temperatures and in equilibrium with its vapor,” *Journal of the American Chemical Society*, vol. 72, no. 7, pp. 2844–2847, 1950.

[6] A. Ladd and L. Woodcock, “Triple-point coexistence properties of the lennard-jones system,” *Chemical Physics Letters*, vol. 51, no. 1, pp. 155–159, 1977.

[7] R. Garcia Fernandez, J. L. Abascal, and C. Vega, “The melting point of ice Ih for common water models calculated from direct coexistence of the solid-liquid interface,” *The Journal of Chemical Physics*, vol. 124, no. 14, p. 144506, 2006.

[8] S. Nosé, “A unified formulation of the constant temperature molecular dynamics methods,” *The Journal of Chemical Physics*, vol. 81, no. 1, pp. 511–519, 1984.

[9] T. Schneider and E. Stoll, “Molecular-dynamics study of a three-dimensional one-component model for distortive phase transitions,” *Physical Review B*, vol. 17, no. 3, p. 1302, 1978.

[10] J. A. Zollweg and G. W. Mulholland, “On the law of the rectilinear diameter,” *The Journal of Chemical Physics*, vol. 57, no. 3, pp. 1021–1025, 1972.

[11] J. S. Rowlinson and B. Widom, *Molecular theory of capillarity.* Courier Corporation, 2013.

[12] A. P. Thompson, H. M. Aktulga, R. Berger, D. S. Bolintineanu, W. M. Brown, P. S. Crozier, P. J. In't Veld, A. Kohlmeyer, S. G. Moore, T. D. Nguyen, *et al.*, “Lammps-a flexible simulation tool for particle-based materials modeling at the atomic, meso, and continuum scales,”

*Computer Physics Communications*, vol. 271, p. 108171, 2022.

[13] H. Yamazaki, M. Takagi, H. Kosako, T. Hirano, and S. H. Yoshimura, "Cell cycle-specific phase separation regulated by protein charge blockiness," *Nature Cell Biology*, vol. 24, no. 5, pp. 625–632, 2022.
